# Unlocking the potential of *Capsicum* Germplasm Collections for Climate Resilience and Fruit Quality

**DOI:** 10.64898/2026.03.25.714358

**Authors:** Anna Halpin-McCormick, Manoj Kumar Nalla, Zachary Radlicz, Andrew Zhang, Nathan Fumia, Tsung-han Lin, Shih-wen Lin, Yen-wei Wang, Herbaud P.F. Zohoungbogbo, Diane Wang, Bryan Runck, Michael A. Gore, Michael Kantar, Derek W. Barchenger

**Affiliations:** Department of Tropical Plant and Soil Sciences, University of Hawaii at Manoa, Honolulu, HI 96822; World Vegetable Center, South and Central Asia Regional Office, Hyderabad, India; Borluag-Ruan International Intern, World Vegetable Center, Tainan, Taiwan; Hawaiʻi Agriculture Research Center, Genetic Resources, Kunia, Hawaiʻi, USA; World Vegetable Center, Tainan, Taiwan; World Vegetable Center, West and Central Africa - Humid and Coastal Regional Office, Cotonou, Benin; GEMS Informatics Center, University of Minnesota–Twin Cities, Saint Paul, Minnesota, USA; Agronomy Department, Purdue University, West Lafayette, Indiana, USA; Plant Breeding and Genetics Section, School of Integrative Plant Science, Cornell University, Ithaca, NY 14853 USA

**Keywords:** *Capsicum*, Climate tolerance, Genomic Prediction, Population improvement, Allele mining, LLM

## Abstract

Climate change increasingly threatens global *Capsicum* (pepper) production. Accelerating the deployment of climate-resilient cultivars requires effective use of genetic diversity conserved in genebanks. We implement a “turbocharging” strategy in *Capsicum* by integrating genome-wide association studies and genomic prediction in a core collection (n = 423), followed by genomic prediction across the global collection (n = 10,250) using the core as a training population. We generated genomic estimated breeding values (GEBVs) for 31 high-accuracy traits (r > 0.5) encompassing hyperspectral phenotypes (heat/control), agronomic performance (heat/control) and fruit quality. To enhance accessibility and decision-making, we developed a large language model (LLM) integrated application that enables flexible, preference-based selection of candidates. By narrowing the parental decision space, this framework streamlines screening of large germplasm collections while balancing climate resilience, quality attributes and market demands. Our approach provides a scalable decision-support system to accelerate climate-resilient *Capsicum* breeding and maximize global genetic resources.

## Main

The world faces interlinked challenges of increasing food insecurity and rising nutritional insecurity, both of which are exacerbated by climate (Tigchelaar et al., 2024; Cooper et al., 2025). Vegetable crops offer a strategic opportunity to address this dual burden. Climate projections indicate that rising temperatures will alter cropping calendars and shift the geographic distribution of suitable production regions (Lee et al., 2018; Marklein et al., 2020; Martínez-Ainsworth et al., 2023). These changes necessitate the selection of genotypes adapted to future environmental conditions while simultaneously enhancing nutritional quality. *Capsicum* provides a compelling model system for examining these intertwined objectives due to its ancient domestication history (Liu et al., 2023), high nutritive value (Kantar et al., 2016) and global culinary ubiquity (Moscone et al., 2007; Paran et al., 2007; Cheng et al., 2016; Wall & Bosland, 1988; DeWitt & Bosland, 1993; Guzman et al., 2011).

Global genebank systems represent the most comprehensive repositories of crop genetic diversity (Byrne et al., 2017). These living collections hold more than seven million accessions across species (Wambugu et al., 2018) and have contributed substantially to cultivar improvement worldwide (Hajjar and Hodgkins, 2007; Dempewolf et al., 2017). Recent advances aim to accelerate the operational use of this diversity by integrating classical strategies such as core collections (Brown, 1989) and focused identification of germplasm strategy (FIGS - Sunitha et al., 2024) with genomic tools including genome wide association studies (GWAS) and genomic selection (GS) (Yu et at., 2016; Li et al., 2018; Yu et al., 2020; Escamilla et al., 2025; Campbell et al., 2025; Halpin-McCormick et al., 2025). While these approaches have substantially improved our ability to characterize genetic resources, challenges remain in efficiently navigating large multidimensional datasets to identify accessions meeting complex breeding objectives. Large language models (LLMs) offer promising complementary tools for engaging searchability and decision support within genomic datasets and their application in plant science is expanding rapidly (Lam et al., 2024; Zhu et al., 2024; Yoosefzadeh-Najafabadi, 2025).

Here we integrate multi-environment phenotyping of the *Capsicum* core collection (n = 423) conducted by the World Vegetable Center (McLeod et al., 2023; Fumia et al., 2023) with genotypic data from the global *Capsicum* germplasm collection (n = 10,250) (Tripodi et al., 2021) to generate genomic estimated breeding values (GEBVs) for agronomic, stress response and nutritional quality traits. These predictions are then rendered searchable through plain language queries (e.g. “*provide me a list of the hottest peppers that are high yielding*”) using an LLM enabled decision support framework.

## Results

### Characterizing the global pepper germplasm collection

Five datasets were used to understand the potential of allele mining for adaptation within the globally available pepper germplasm. In this study a subset of the *Capsicum* core collection (n = 280/423 accessions, n = 340,734 SNPs) (hereafter core collection) were phenotyped for 73 traits (**Table S1**). This genotyped core collection (**Table S2**) were phenotyped under a control condition (21.4 ± 3.82°C), heat stress-1 (HS1 = 28.8 ± 1.42°C) and heat stress-2 (HS2 = 28.1 ± 2.50°C) conditions (**Figure S1**). A subset of the core collection (n = 329 accessions) had been previously phenotyped for quality (n = 23 traits) (**Table S3**) (McLeod et al., 2023). Finally, GEBVs for the entire *Capsicum* germplasm collection (hereafter global collection) (n = 10,250 accessions, n = 23,462 SNPs) were developed using a shared n = 259 accession overlap with the core collection (**Table 4**). We first recapitulated the population structure of the core collection (n = 340,734 SNPs) using a principal component analysis (PCA), hierarchical clustering, and fastSTRUCTURE analysis (**Figure S2-6**). There was clear separation among species and domestication status (**Figure S2-6**). There was also clear admixture within the *Capsicum annuum* accessions, indicating that there likely has been considerable germplasm exchange over time. The same set of analyses were conducted in the global germplasm collection (n = 10,250 accessions, n = 23,462 SNPs), where we observed similar clustering with clear separation among species and some evidence of admixture (**Figure S5-7**).

### Understanding stress tolerance through GWAS in the *Capsicum* core collection

Best Linear Unbiased Estimates (BLUEs) were estimated from phenotypic data within each dataset. A stress tolerance index (STI) was subsequently calculated using the adjusted genotype means, thereby reducing environment specific variation (e.g. confounding variation and error variance) (Sadrarhami et al., 2010). An STI normalizes across the stress environments and thus can identify SNPs that are associated with stable performance across environments. To dissect the genetic basis of agronomic performance and stability under thermal stress, genome-wide association studies (GWAS) were conducted using four models (BLINK, MLMM, MLM with PC + K, and FarmCPU) on 73 traits. STI GWAS was conducted with 340,734 SNPs on 280 genotypes (*Capsicum annuum* only) from the core collection. The agronomic traits assessed in this study were grouped into eight broad functional categories for downstream interpretation (**Table S1**); canopy spectral properties (n = 19; green reflectance, hue, and NDVI indices), physiological traits (n = 24; leaf temperature, leaf angle, light interception, and leaf inclination), leaf area and canopy architecture (n = 10), vegetative growth (n = 8; plant height, biomass, and growth rate), yield components (n = 5; fruit number, weight, length, width, and shape index), phenological timing (n = 4; days to anthesis and maturity after sowing and transplanting), pollen traits (n = 2; pollen concentration and activity), and direct yield (n = 1; fruit yield in tonnes per hectare) (**Table S1, Figure 1**).

**Figure 1.**
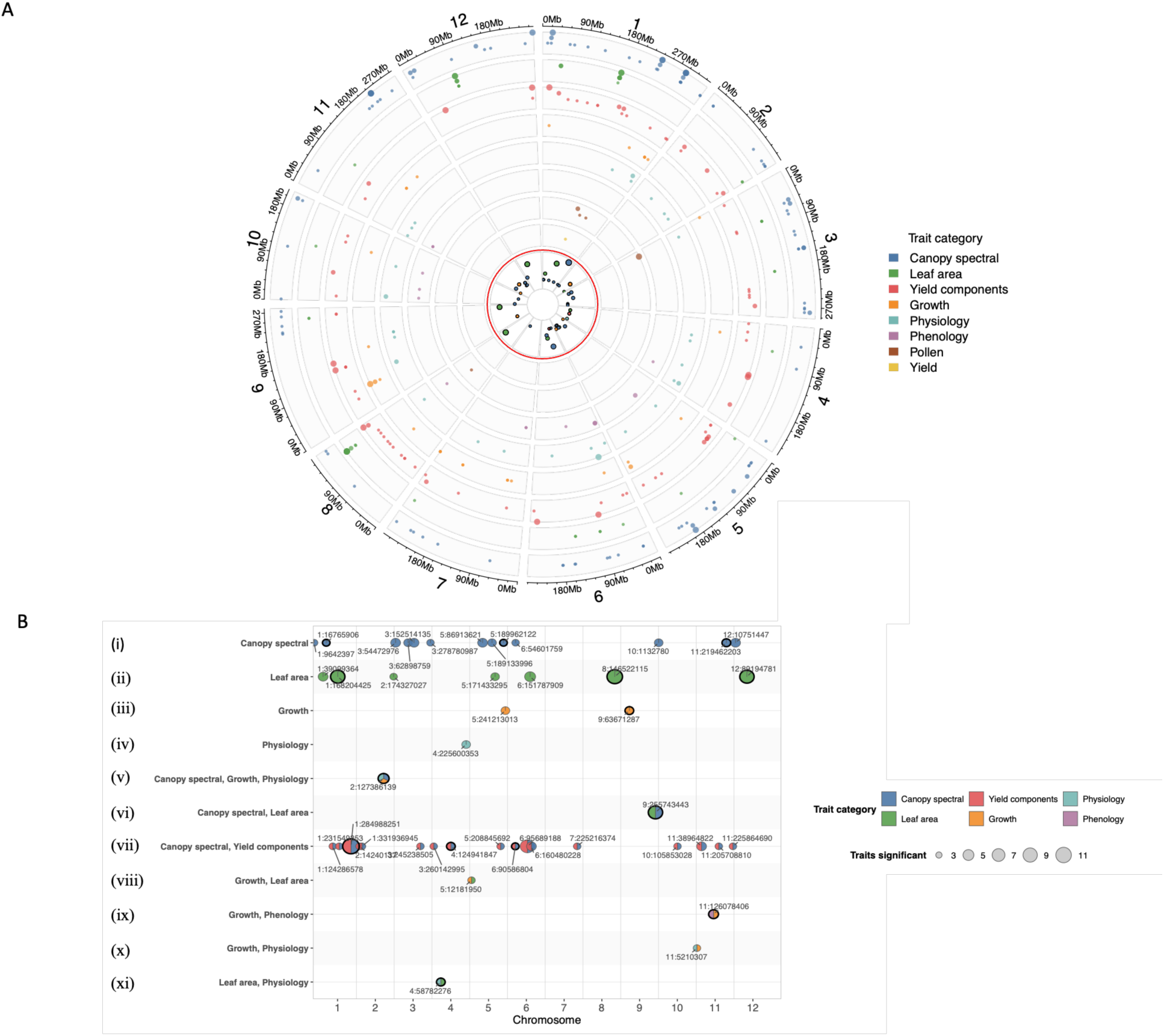
Genome-wide association study (GWAS) of stress tolerance index (STI) across eight functional trait categories in the *Capsicum annuum* core collection. GWAS was performed on 280 *C. annuum* accessions from the core collection using four models (BLINK, MLMM, MLM with PC + K, and FarmCPU) and 340,734 SNPs across 73 traits. Agronomic traits were grouped into eight functional categories: canopy spectral properties (blue), leaf area (green), yield components (red), vegetative growth (orange), physiology (teal), phenology (purple), pollen (brown), and direct yield (yellow). (**A**) Chromosomal distribution of 325 SNPs significant in ≥2 GWAS models after Benjamini-Hochberg false discovery rate correction (FDR < 0.05). Point size reflects the number of models in which the SNP reached significance (2, 3, or 4 models). Inner track (in red) shows the chromosomal distribution of 46 high-confidence loci (**B**) Chromosomal distribution of high-confidence loci (n = 56). Circles represent SNPs identified at the intersection of multi-model replication (≥3 GWAS models) and cross-trait pleiotropy (≥3 traits), with point size reflecting the number of associated traits. SNPs outlined in black were significant in all four GWAS models. SNP identifiers (Chr:Position) are labelled for each locus.

There were 325 significant SNPs identified in ≥ 2 GWAS models (FDR < 0.05) across all traits (**Table S5, Figure 1A**). Separately, 264 SNPs were identified as significant and pleiotropic (observed in ≥ 3 traits) (**Table S6**). SNPs associated with pollen and yield traits were included here despite not meeting the pleiotropy threshold, as only two pollen traits were measured (pollen concentration and pollen activity), and fruit yield was a single trait, making it impossible for any SNP to reach the ≥3 trait threshold by definition. Intersecting these two sets identified 46 high confidence SNPs that were observed to be pleiotropic (≥ 3 traits) and present in ≥3 GWAS models (**Table S7, Figure 1B**). LD decay patterns in the core collection (**Figure S23**) provide additional context for interpreting significant SNPs detected. Given the observed LD decay distance, r^2^ = 0.2 at approximately 56kb (**Figure S23**), significant SNPs are expected to tag causal variants within ∼50-60 kb intervals.

The 46 high confidence SNPs spanned five broad phenotypic domains. The largest group (n = 17) comprised SNPs associated with both canopy spectral indices and fruit number, where huebin0 and huebin5 were consistently co-detected with fruit number across multiple chromosomes (**Figure 1B, vii**). A further 13 SNPs were associated exclusively with canopy spectral traits (**Figure 1B, i**). Ten SNPs were associated with leaf area (7 exclusively, **Figure 1B, ii**), including large-effect loci at Chr1:168,204,425 and Chr8:146,522,115, each detected across 9-10 leaf area traits and all four GWAS models. The remaining SNPs spanned vegetative growth (n = 5, including height, growth rate, and biomass), physiology (n = 1, leaf inclination and light penetration) and a notable pleiotropic locus at Chr2:127,386,139 associated with biomass, growth rate, height, light penetration, and canopy spectral traits across three trait categories (**Table S7, v**). These results suggest that canopy spectral properties and leaf area represent an adaptive strategy underpinning STI variation in the core collection (**Figure 1B**).

Principal Component (PC) adjusted percent variance explained (PVE) was estimated for each SNP x trait combination (**Table S8)**, with per SNP summaries (mean, minimum and maximum PVE across traits) reported in **Table S7**. The co-detection of loci for canopy spectral indices and fruit number reflects residual population stratification in fruit number STI, rather than genuine pleiotropy, as PC adjusted percent variance explained (PVE) for fruit number was near zero at these loci (e.g. Chr1:124286578, mean fruitno PVE = 0.16%), while the same SNPs explained 11.6-18.5% of huebin0 and huebin5 STI variance respectively (**Table S8**). Three loci showed independently significant fruit number effects Chr6:90,586,804 (fruitno PVE = 43.1%), Chr4:124,941,847 (19.8%) and Chr3:245,238,505 (15.7%) and represent candidate loci with potential direct roles in fruit number output under heat stress.

### Environment specific genomic prediction and rank changes

Heat tolerance exhibits quantitative genetic inheritance in many plant systems (Yang et al., 2002; Healy et al., 2018; Zhang et al., 2022). Therefore we sought to identify lines potentially associated with heat responsiveness. Genomic estimated breeding values (GEBVs) were calculated using 340,734 SNPs in the core collection (n = 423) for 73 traits (**Table S9-11**) for the three different conditions, control, HS1 and HS2 (**Figure S12A, Figure S8-11**) and GEBV mean distributions were compared (**Figure S12B**). Thirteen of the 73 traits (17.8%) had a prediction accuracy higher than 50% across all three environments (biomass final plant, biomass final plot, fruit length, fruit number, fruit shape index, fruit weight, fruit width, leaf angle daily, leaf area daily, leaf area daily plant, leaf area projected daily, leaf area projected plant, yield - **Figure S12B**, **Figure S8-11**). A threshold of 0.5 has been empirically determined to be reasonable for high accuracy in several different crop systems (Heslot et al., 2012; Crossa et al., 2014; Desta and Ortiz, 2014; Schrauf et al., 2021). This also acts as a heritability filter, where we see this prediction accuracy corresponds to mid and high heritabilities (**Figure S22).**

To identify the most pronounced rank shifts, the top 20% of genotypes with largest rank change between conditions were retained for additional analysis **(Figure S13, Table S12-13**). Accessions with reduced performance (lower GEBVs) under heat stress may be classified as control-responsive, while those with improved performance (higher GEBVs) under heat may be considered heat-responsive (**Figure S13**). Among these, 28 accessions displayed high breeding values under heat stress and were considered heat tolerant (**Figure S13A**), while 57 accessions showed superior performance under control conditions (**Figure S13B**) (**Table S12-13**). Distinct rank reordering of accessions across conditions (**Figure S11 & S13, Table S9-11**) indicated that accession utility is environment dependent, with some genotypes performing consistently across heat conditions (**Figure S13A**) and others across cooler control conditions (**Figure S13B**) and exhibiting strong condition specific performance.

Following this analysis on the core collection, GEBVs were calculated for the global collection using the core collection as the training set (n = 259 overlapping accessions) (**Table S4, Table S14-16**). While this was an appreciably small training set (2.5%), prediction accuracy was relatively high for many of the phenotypes (**Figure S14-16**). There were 17 traits with prediction accuracies consistently above the r > 0.5 threshold (**Figure S17**). In the global germplasm collection, there was variation in GEBV distributions among the stresses, where some agronomic traits in the global collection had distributions that exceeded the core and in other incidences the distributions were similar (**Figure S21**). In addition to mean GEBV shifts, the HS1 and HS2 distributions exhibited pronounced changes in spread relative to control. Notably some traits displayed bimodal distributions, suggesting potential sub population structure or divergent responses among accessions (**Figure S12B, Figure S20B, Figure S21**).

### Stress tolerance genomic prediction across the core and global collections

To further characterise the genomic basis of heat stress tolerance, stress tolerance indices (STI) derived from environment specific BLUEs were used as the phenotypic input for genomic selection across the 72 agronomic traits, with GEBVs calculated for the core collection and the global collection (**Figure 2, Table S17-18**). Prediction accuracies varied across traits and between collections with yield (core PA = 0.59, global PA = 0.56), yield components (core PA = 0.64, global PA = 0.66), leaf area (PA = 0.63 both), and growth traits (core PA = 0.55, global PA = 0.63) exceeding the r > 0.5 threshold in both collections, while physiology traits showed consistently lower accuracies (core PA = 0.28, global PA = 0.36) (**Figure 2, Figure S24 and S25**). Canopy spectral and yield component distributions were broadly similar between collections, suggesting the core adequately captures the range of canopy and yield component STI variation. Notably, yield STI alone showed a contrasting pattern, with the core collection shifted left relative to the global collection.This divergence, combined with modest prediction accuracy (PAs 0.59 and 0.56) indicates that the global collection may require targeted phenotyping to identify rare high-yield-STI accessions that fall outside the distribution captured by the core training set (**Figure 2**).

**Figure 2.**
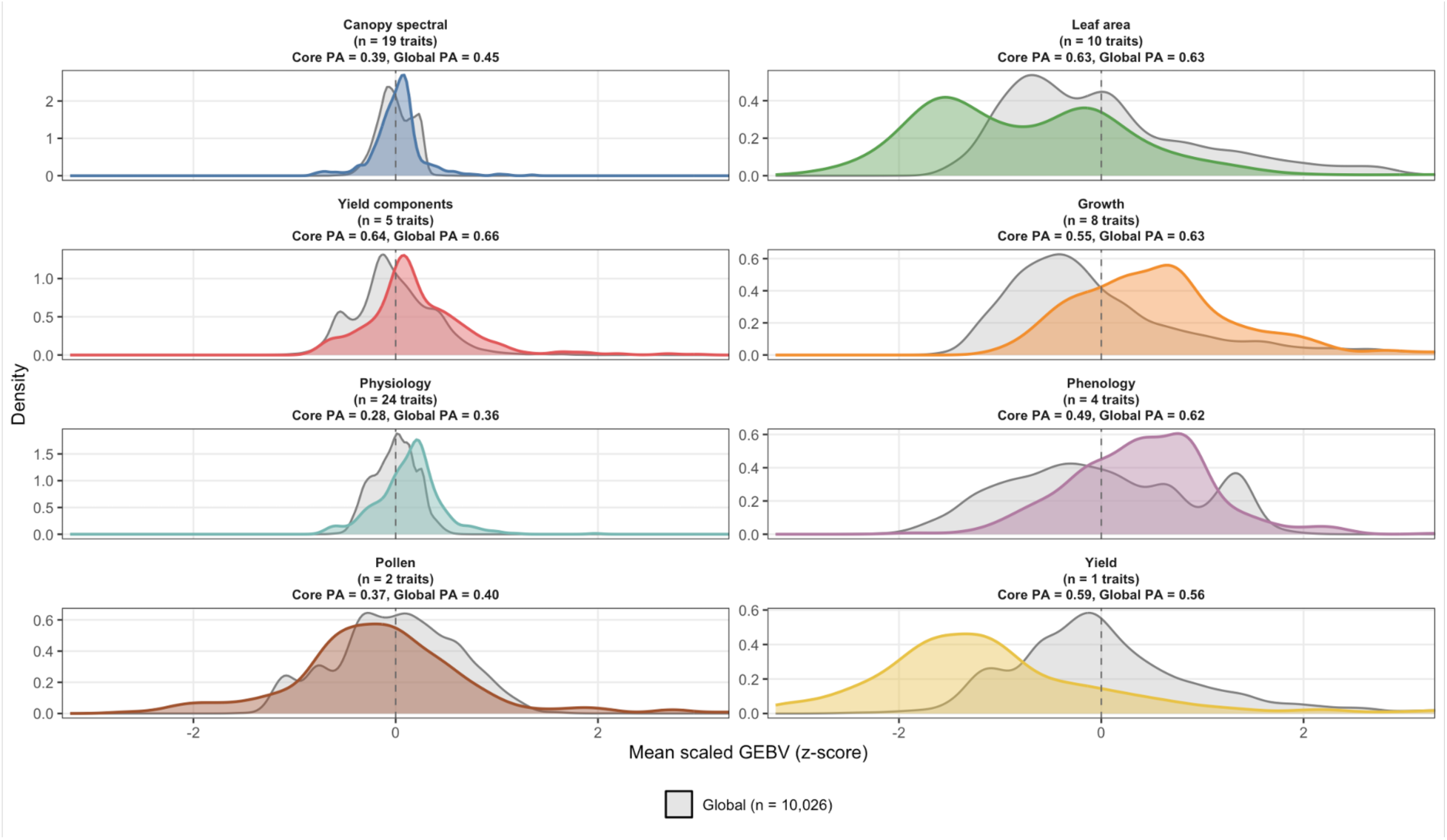
Distributions of mean scaled genomic estimated breeding values (GEBVs, z-scores) across eight trait categories for the core (n = 423) and global (n = 10,026) *Capsicum annuum* collections. GEBVs were estimated using rrBLUP with the stress tolerance index (STI) as the response variable. The core collection model was trained on 150 phenotyped lines, while global collection GEBVs were predicted using a model trained on all 259 phenotyped lines common to both datasets. Prior to averaging across traits within each category, GEBVs were z-score scaled within each trait across both collections combined, allowing direct comparison of breeding value distributions. Mean prediction accuracy (PA) per category represents the average cross-validated PA (10-fold, 50 replicates) across all traits within that group, estimated independently for each collection. Core collection distributions are shown in colour, while global collection distributions are shown in grey.

### Genomic prediction of market quality traits

*Capsicum* species have multiple purposes in global agriculture and a multitude of markets that differ in their breeding priorities. Bell pepper markets typically value traits such as yield, fruit size, shape, colour and shelf life. While chili pepper markets prioritize host resistance to pests and disease, yield, dry matter, carotenoid content and pungency or heat level. These differences reflect diverse regional culinary preferences (McLeod et al., 2023). Using data from McLeod et al., GEBVs were calculated for 23 quality traits (**Figure 3, Table S3**) using a training set of n = 150 accessions for the core collection with 18/23 traits (78.2%) having prediction accuracies of r > 0.5 (**Figure S20B**, **Figure S18, Table S19**). Similarly, GEBVs were calculated for quality traits for the global *Capsicum* collection using a training set of n = 329 accessions (**Figure S20B**, **Table S20**) with 17/23 (73.9%) having prediction accuracies of r > 0.5 (**Figure S19, Table S20**) offering insights into the potential of improving quality using the global collection (n = 10,250).

**Figure 3.**
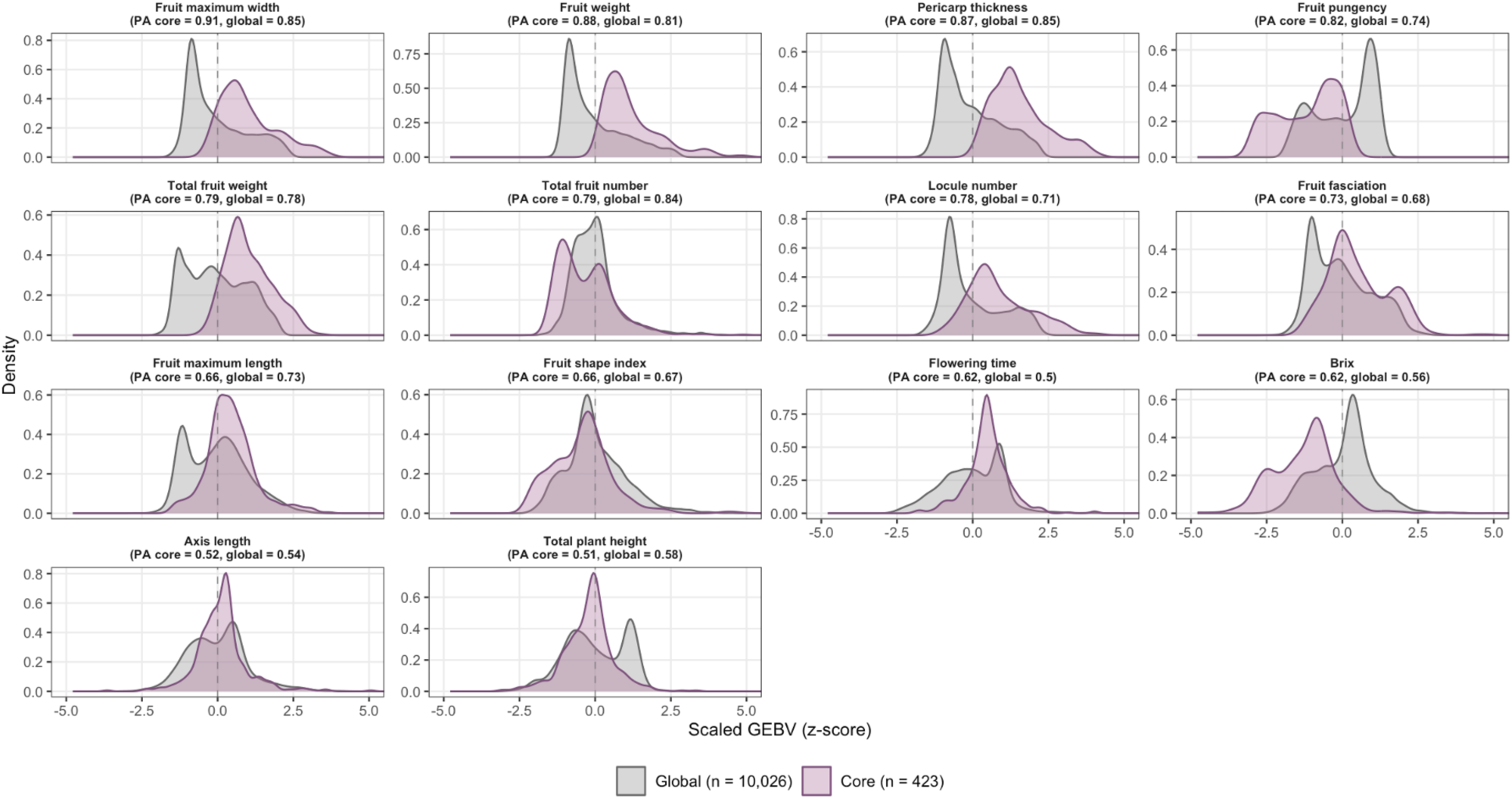
Distributions of scaled genomic estimated breeding values (GEBVs, z-scores) for quality traits in the core (n = 423) and global (n = 10,026) *Capsicum annuum* collections. GEBVs were estimated using rrBLUP with phenotypic data from the McLeod et al. quality trait dataset. The core collection model was trained on 150 phenotyped lines, while global collection GEBVs were predicted using a model trained on all 329 phenotyped lines common to both datasets. Prior to plotting, GEBVs were z-score scaled within each trait across both collections combined, allowing direct comparison of breeding value distributions. Prediction accuracy (PA) reported per trait represents the cross-validated PA (10-fold, 50 replicates) estimated independently for the core and global collections. Traits are ordered by decreasing PA in the core collection. Purple distributions represent the core collection, grey distributions represent the global collection.

Trait specific differences were observed in GEBV distributions for quality between the core and global collections (**Figure 3, Figure S20B**) (**Table S19-20**). Notably, mean values for brix, fruit, pungency and plant height were higher in the global collection, indicating that selection from the larger collection may provide greater opportunity to maximize benefits from these traits (**Figure 3, Figure S20B)**.

### Overlap between climate resilience and quality

By integrating GEBVs across abiotic stress and quality traits, accessions can be identified that are suitable for diverse market classes. There were clear patterns with some quality traits being positively correlated with increased heat stress and others being negatively correlated with heat stress (**Figure S20A)**. It is possible to therefore leverage GEBVs to identify parents for diverse market classes, by setting specific thresholds for relevant traits for each specific uses, subsetting the total collection to only those accessions that pass filtering levels. Trait distributions differed among stress levels (**Figure S20B**, **Figure S21**), indicating that there may be unmined potential in the global collection for some traits that is not well represented in the core collection. This also shows how common variation in a core collection can be leveraged to identify rare variation in a global collection.

### LLM search tool to enable parental selection

Using a thresholding approach to sort accessions by breeding values requires a large amount of species and market knowledge. This can limit accessibility and utility of the *Capsicum* collections. To overcome this, we developed a plain language LLM-assisted search tool that allows users to subset accessions using natural language queries in addition to traditional trait thresholds (**Figure 4**). The application integrates GEBVs derived from both STI based genomic selection across 73 agronomic traits (**Table S17/18**) and quality trait (**Table S19/20**) predictions to enable simultaneous querying. The GEBVs for traits of interest are searched and a subset of accessions is returned which match the users criteria. The LLM uses programmatic tools exposed through an MCP (Model Context Protocol) server to convert the filtering plan into frontend application, through slider adjustments and reading. This allows the user to see the effects of the search tool updating live on the application and to make changes to recommended accessions. To enable multi-trait selection, we implemented both a simple weighted index and a prediction accuracy adjusted Smith-Hazel genomic selection index within the GEBV Explorer application.

**Figure 4.**
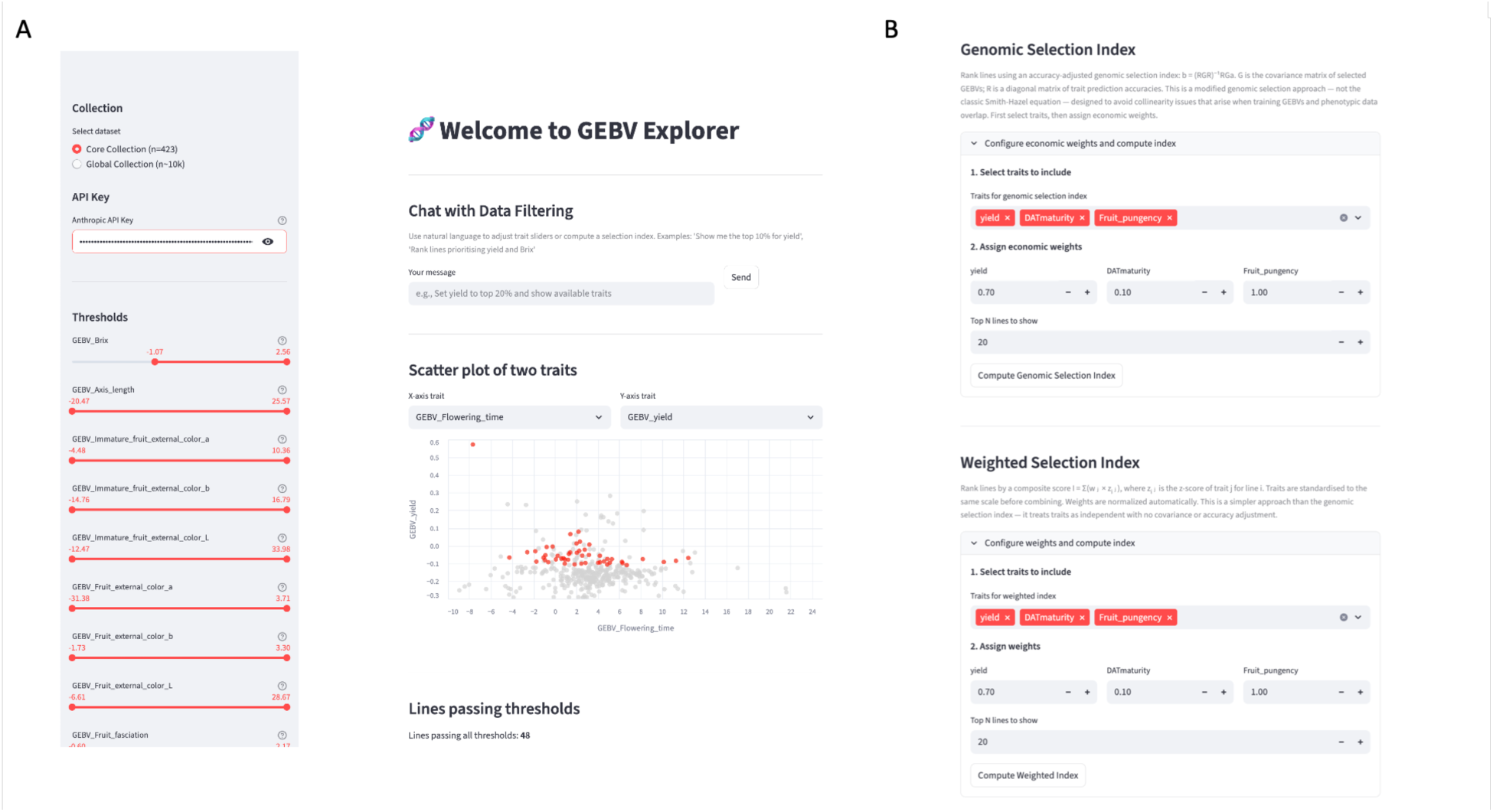
LLM-integrated genomic estimated breeding value (GEBV) application for interactive selection within *Capsicum* germplasm. The application enables user driven exploration of genomic prediction outputs across 23 quality and 73 agronomic traits. (**A**) The main interface supports direct GEBV threshold filtering via interactive sliders and natural language queries (personal Claude API key required), allowing users to subset lines based on trait specific criteria. A dynamic scatter plot visualizes relationships between selected traits and filtered candidate lines are displayed with export functionality. Results for the *Capsicum* core and global collections are available for searching. **(B**) Two index-based selection approaches are also available: (i) a weighted selection index, where standardized trait values are combined using user-defined weights and (ii) a genomic selection index, incorporating trait covariance structure and prediction accuracy (Smith-Hazel framework) with user specified economic weights. These tools enable flexible, multi-trait optimization and rapid identification of superior lines for breeding decisions.

## Discussion

While genebanks represent vast reservoirs of allelic diversity, their operational deployment in breeding programs remains restricted due to difficulties in evaluating and prioritizing accessions at scale (Hajjar and Hodgkins, 2007; Dempewolf et al., 2017; Wambugu et al., 2018; Campbell et al., 2025). Here, the generation of genomic estimated breeding values (GEBVs) for agronomic, stress and quality traits quantitatively ranks accessions and enables users to easily prioritize candidate breeding lines for multiple potential breeding objectives **(Figure 2**, **Figure 3, Figure S13, Figure S20, Table S9-S11, S14-S20**). By coupling genomic prediction with natural language filtering and trait thresholding, we reduce barriers associated with multi-trait selection, operationalizing genebank diversity through predictive decision support. Substantial investment has been directed toward genotyping and phenotyping global germplasm collections (Langridge & Fleury., 2011; McCouch et al., 2013). However, declining genotype costs relative to large-scale multi-environment phenotyping creates a strategic opportunity to leverage phenotype core collections as training populations for genomic prediction across entire collections (Yu et al., 2016). This approach enables the projection of trait values onto thousands of unphenotyped accessions, expanding the actionable diversity available to breeders.

Pepper has a large number of market classes (n = 88; Ambali et al., 2025), each defined by unique combinations of agronomic and horticultural traits. Consequently, breeders require selection tools adaptable to diverse breeding objectives. The integration of genomic prediction with natural language based filtering allows complex trait combinations to be queried efficiently. For example, user defined search criteria such as “*provide me accessions that show high pungency, high yield and early maturing*” returns the following 8 accessions from the core collection GPC001360, GPC003150, GPC029990 GPC049590, GPC076940, GPC093310, GPC113230 and GPC123700.

These accessions constitute a candidate parent set for crossing with locally adapted material to generate hybrids that combine regionally important traits with globally relevant adaptive alleles. These results indicate that targeted market segments will require strategic utilization of the genetic diversity present within collections, with identification of optimal parental combinations depending on integrating knowledge of local consumer preferences with underlying trait correlations.

This class of methods termed ‘germplasm genomics’ enables multiple pathways for crop improvement with selection depending on underlying genetic architecture (Wambugu et al., 2018; Sanchez et al., 2023). Traits governed by major effect loci are amenable to marker assisted selection (MAS) (**Figure 1)**, whereas traits with polygenic control are better suited to genomic selection (GS) (**Figure 2 and 3**). Both improvement strategies emerge from the same multi-environment screening, demonstrating that genebank based evaluations can simultaneously support allele mining for discrete loci and population improvement for complex traits.

As climate instability reshapes agricultural production landscapes, predictive deployment of global genetic resources will become increasingly central to food system resilience and nutritional security (Khaitov et al., 2019; Tigchelaar et al., 2021; Cooper et al., 2021; Ortiz-Bobea et al., 2021). This is particularly relevant in vegetable crops such as *Capsicum,* where micronutrient content and phytochemical composition contribute to dietary diversity (Olatunji & Afolayan, 2018). The pronounced rank reordering of accessions across different thermal environments demonstrates the vulnerability of accessions to climate variability. Accessions that perform strongly under control conditions frequently decline under heat stress, with distinct subsets of accessions improving in relative ranking between experimental timepoints (**Figure S13A/B**). This dynamic suggests that climate change may substantially affect breeding priorities and that predictive tools capable of environment specific ranking are important for the continued development of climate resilient food systems (Challinor et al., 2014). Additionally, this reflects a broader tension between selecting generalist genotypes that maintain stable performance across environments and specialist genotypes optimized for specific climatic conditions (Levins 1962; Finlay & Wilkinson., 1963; Ewing et al., 2019). Variation in GEBVs across abiotic stress and quality traits reveals accessions that simultaneously exhibit climate resilience and favorable market attributes, indicating that these characteristics are not inherently antagonistic in *Capsicum* (**Figure S20A)**. This creates an opportunity to co-optimize yield stability and quality across diverse market preferences.

## Conclusion

By integrating genomic predication and natural language decision support tools, we demonstrate a framework enabling global germplasm collections to function as predictive breeding resources. This approach reduces the dimensionality of accession selection and supports the identification of parental lines with stable performance under stress while considering market driven quality objectives. Through Standard Material Transfer Agreement (SMTA), this resource has the potential to accelerate improvement by enabling breeders to quickly assess the global *Capsicum* collection for their desired trait combinations of interest. This approach is transferable to other crop genebank collections where appropriate genotypic and phenotypic data has been collected, facilitating targeted selection of parental lines from large collections.

## Materials and Methods

### Plant Material

A core collection for chili pepper has previously been defined (McLeod et al., 2023) and was utilized in this study. This core (n = 423) consists of *Capsicum annuum* (n = 397), *Capsicum baccatum* (n = 7), *Capsicum chacoense* (n = 1), *Capsicum chinense* (n = 12) and *Capsicum frutescens* (n = 6) accessions (**Table S1**). The global collection (n = 10,250) for *Capsicum* has also been previously defined (Tripodi et al., 2021) and was additionally utilized in this study.

### Phenotypic Trials

Entries were evaluated during three seasons as previously described in Fumia et al., 2023. Briefly, two heat stress seasons characterized by long periods of stable, but high temperatures and one cool season (control season) were observed during the study period. All three field trial experiments were conducted at the World Vegetable Center in Shanhua, Tainan, Taiwan (lat. 23.1 ° N, long. 120.3 ° E, elevation 12 m.a.s.l.). In total, 73 phenotypic observations (Table S1), including plant morphology, color, leaf temperature, pollen quality and yield component data, were recorded. Automated data were collected using the PlantEye 3D-Spectral Scan F500 (Phenospex, Heerlen, the Netherlands). Manual data (classical phenotypes) included days to flowering, fruit maturity, yield components of fruit length, width, weight, and yield, as described by Barchenger et al., (2018, 2020) as well as pollen concentration and activity using impedance flow cytometry following the protocol of Lin et al. (2022). Best linear unbiased estimates (BLUEs) were calculated separately within each environment (Control, HS1, HS2) using a linear mixed model with accession as a fixed effect and replicate as a random effect. BLUEs were estimated using the lme4 package in R, with marginal means extracted using emmeans.

### Sequencing

Whole genome shotgun was performed on 423 *Capsicum* ssp. accessions which constitute the G2P-SOL pepper core collection (McLeod et al., 2023) and capture global pepper diversity. Alignment to the *Capsicum annuum* var. *annuum* (pimento, cultivar: Zhangshugang) reference genome (GCA_030867735.1) and subsequent SNP calling using the GATK (v4.5.0.0) (DePristo et al., 2011) pipeline resulted in 423,926,693 SNPs (raw_SNP_all.vcf.gz). For filtering, variants were filtered using bcftools (v1.11) (Danecek et al., 2021) and VCFtools (v0.1.16) (Danecek et al., 2011) and included the following criteria (i) restriction to biallelic variants (--max-alleles 2), (ii) setting a minimum read depth (DP) between 4 and 50 (--minDP 4, --maxDP 50), (iii) filtering out genotypes with a genotype quality (GQ) below 10 (--minGQ 10), (iv) excluding variants with a quality score (QUAL) below 30 (--minQ 30), (v) removing variants with a minor allele frequency (MAF) below 0.05 (--maf 0.05) and (vi) retaining variants only if present in at least 90% of samples (--max-missing 0.9). This resulted in 44,786,599 SNPs. Due to the presence of unplaced contigs, the vcf was partitioned to only the 12 chromosomes of *Capsicum annuum* and resulted in 44,783,415 SNPs. Linkage disequlibirium (LD) pruning was performed using PLINK (v1.90b6.21) (Purcell et al., 2007) with a sliding window size of 5000 SNPs, a step size of 500 SNPs and a *r^2^* threshold of 0.2. This resulted in 340,734 variants remaining.

### Population Structure

The above dataset was examined for population structure using “SNPRelate” For the Principal Component Analysis (PCA) and Hierarchical Clustering of Principal Components (HCPC) using “FactoMineR”, using only bi-allelic SNPs further filtered for linkage disequilibrium (0.2).

### Estimation of phenotypic BLUEs

Phenotypic data were collected in a randomized complete block design comprising three environments, three replicates per environment (control, heat stress-1 and heat stress-2) for 280 accessions. Best linear unbiased estimates (BLUEs) were obtained for each trait using linear mixed-effects models implemented in the lme4 package (Bates et al., 2015) in R. For across environment BLUE estimation, phenotypic values were modelled with accession as a fixed effect and environment and replicate nested within environment as random effects, according the the following model:

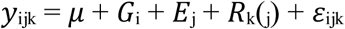

*y*_ijk_ is the observed phenotypic value of accession *i* in environment *j* and replicate *k*

μ is the overall mean

*G*_i_ is the fixed effect of accession *i*

*E*_j_ is the random effect of environment *j*

*R*_k(j)_ is the random effect of replicate *k* nested within environment *j*

ε_ijk_ is the residual error term

Accession level BLUEs were extracted using estimated marginal means (emmeans) package (Lenth et al., 2025) and used as adjusted phenotypic inputs for downstream analyses. In addition, environment-specific BLUEs were also estimated separately for each environment using models with accession as a fixed effect and replicate as a random effect.

### STI Genome Wide Association Study (GWAS)

Environment specific genotype BLUEs (control, heat stress-1, heat stress-2) were used to compute a Stress Tolerance Index (STI) for genome-wide association analyses using the dplyr package in R (Wickham et al., 2023), based on an extension of the stress tolerance framework proposed by Fernandez (1992), integrating genotype performance across stress and non-stress environments.

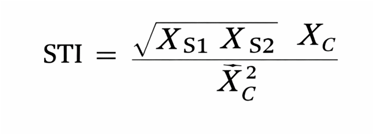

Where

X_S1_ = Trait value under stress condition 1(HS1)

X_S2_ = Trait value under stress condition 2 (HS2)

X_C_ = Trait value under standard commercial production practices (control)

X^-^_c_ = mean trait value under standard commercial production practices

The STI was used to identify genotypes capable of maintaining high performance under both heat stress and control conditions. A high STI value indicates a greater degree of heat stress tolerance and phenotypic stability. A genome wide association study was conducted to identify genetic loci associated with agronomic traits (**Table S1**) for the 280 *Capsicum annuum* accessions which had phenotype data. Analysis was performed using the GAPIT (v3) package (Wang and Zhang, 2021) implementing the FarmCPU, Blink, MLMM and MLM (PC 1-5 + K) models. To account for multiple testing, marker-trait association p-values were adjusted using the Benjamini-Hochberg false discovery rate (FDR) correction. Associations with an FDR-adjusted P < 0.05 were considered statistically significant. Percent variance explained (PVE) was estimated for each high-confidence SNP × trait combination as the incremental r^2^ attributable to SNP dosage after partialling out population structure, calculated as r^2^ (PC1-5 + SNP) – r^2^(PC1-5 only) (**Table S7/8**).

### Genomic Selection (GS)

Three genomic prediction methods were examined in all cases namely: (i) RR-BLUP, (ii) G-BLUP with an exponential kernel and (iii) G-BLUP with a Gaussian kernel. For model evaluation, 10-fold cross validation was repeated 50 times (k = 10, reps = 50) on the training set for each variable. Genomic selection for heat tolerance (control, heat stress-1 and heat stress-2) was performed for the core chili collection (n = 423) using 340,734 SNPs and 73 phenotypic traits (**Table S1**) collected at the World Vegetable Center in Taiwan. The training set consisted of 150 samples which were selected randomly from the collection (**Table S2**) with the test set consisting of the remaining 273 samples.

Genomic selection for heat tolerance (control, heat stress-1 and heat stress-2) was similarly performed for the global chili collection (n = 10,250) using 23,462 SNPs and 73 phenotypic traits (**Table S2**). The training set consisted of 280 samples (**Table S4, S17**) with the test set consisting of the remaining 9,970 samples. Genotype data for the global pepper collection was sourced from Tripodi *et al*. 2021 with 5.92% missingness which was imputed using the mean for each NA marker. GEBVs for the 73 traits were calculated similarly. There were 224 control or replicate samples of *C. annuum* cv. “CM334” in the Tripodi vcf file. These were removed from the final tables (**Table S14, S15, S16, S18**) leaving 10,026 samples.

Genomic selection was additionally performed for 23 quality and traditional traits (**Table S3**). This phenotype data was sourced from McLeod et al. 2023 with 329/350 sample overlaps (**Table S2/S4**) with both the core and global collections which were available for use as the training set. For the GS for quality for the core collection, the same 150 lines used previously as the training set were kept for training (**Table S2**). For the GS for quality for the global collection, all 329 lines were used as training leaving 9,921 samples as the test set.

### Relationship between Heritability and Prediction Accuracy

Genotypic data were obtained from a filtered and linkage disequilibrium pruned VCF file. Genotypes were extracted using the vcfR package (v1.15.0) (Knaus & Grünwald, 2017) and converted from genotype strings into numeric allele dosages. SNPs were filtered with Minor allele frequency (MAF) > 0.05 and missing data of < 10%. Missing genotype values were mean-imputed for each marker, on a per marker basis and a genomic relationship matrix was calculated. Narrow sense heritability (h^h^) was estimated for each trait within each condition using a genomic best linear unbiased prediction (GBLUP) model implemented in the sommer package (v4.4.4) (Covarrubias-Pazaran, 2016) (**Figure S19**).

### Linkage Disequilibrium (LD) Decay

LD decay was estimated using the *Capsicum annuum* filtered SNP dataset. Pairwise LD (r^2^) was calculated between SNPs within chromosomes and restricted to pairs separated by ≤ 1 Mb. Pairwise distances were grouped into physical distance bins (0-1 Mb) and mean r^2^ was calculated per bin. An exponential nonlinear regression model was fitted to the genome wide mean r^2^ values to estimate the LD decay curve. Half decay distance and the distance at which r^2^ reached 0.2 were derived from the fitted model (**Figure S23**).

### Correlation Analysis of Phenotype x Environment Combinations

Pearson correlations were calculated among all shared phenotypic traits across control, heat stress-1 and heat stress-2 conditions after filtering variables with > 50 % missing data, fewer than 10 non-missing observations or zero variance. Correlations were calculated using pairwise complete observations and visualized without clustering or reordering (219 × 219 matrix) with fixed scale −1 to +1 (**Figure S24**).

### Design of a LLM Search Tool

GEBVs derived from STI-based genomic selection models for the *Capsicum* core collection (n = 423) and for the *Capsicum* global collection (n = 10,026) for 23 quality and 73 agronomic traits were used as inputs for the application. The application was implemented in Python using a Streamlit interface connected to a lightweight API server. Each trait was dynamically converted into a percentile based threshold slider. The integrated LLM converts user queries into a JSON structured filtering plan that specifies the columns to filter and sort, while accommodating trait synonyms and metadata. To enable natural language querying of multi-trait combinations and to allow an external LLM agent to manipulate trait thresholds through API calls, a Model Context Protocol (MCP) server was implemented, with this architecture enabling human driven user interface filtering. To facilitate multi-trait selection, two genomic selection indices were implemented in the GEBV Explorer. First, a weighted linear index was calculated as

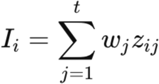

where

*z_ij_* = z-scored GEBV for line i, trait j

*w_j_* = economic weight

I_i_ = composite index score

*t* = number of traits included in the index

Second, a prediction accuracy adjusted Smith-Hazel genomic selection index was implemented and was calculated as

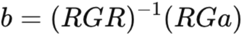

where

*b* = vector of index coefficients applied to each trait

*G* = covariance matrix among the selected GEBV traits

*R* = diagonal matrix containing genomic prediction accuracies for each trait

*a* = vector of user-defined economic weights

The resulting coefficient *b* accounts for genetic covariance among traits as well as differences in genomic prediction accuracy. The final index score for each genotype was then calculated as

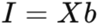

where

*I* = genomic selection index score

*X* = matrix of GEBVs for the selected traits

*b* = vector of derived index coefficients.

This application is available at the following link for the *Capsicum* core and global collections (https://github.com/GEBV-collab/GEBV_Explorer_V2.git). A personal Claude API key is required for interaction with the chat feature.

## Acknowledgements

We acknowledge long-term strategic donors to the World Vegetable Center: Taiwan, United Kingdom, United States, Australia, Germany, Thailand, Philippines, South Korea and Japan. Ministry of Agriculture of Taiwan funded the current Genomic Selection work. Funding for this research was provided by the Ministry of Agriculture of Taiwan (project number 114AS-1.2.3-AS-02), the European Union’s Horizon 2020 Research and Innovation Programme under Grant Agreement No. 677379 (G2P-SOL project: Linking genetic resources, genomes and phenotypes of Solanaceous crops), the Rural Development Administration of Korea as well as the long-term strategic donors to the World Vegetable Center: Taiwan, UK aid from the UK Government, United States Agency for International Development (USAID), Australian Centre for International Agricultural Research (ACIAR), Germany, Thailand, Philippines, Korea, and Japan. Additional funding provided by The College of Tropical Agriculture and Human Resources, University of Hawaii at Manoa; USDA Cooperative State Research, Education and Extension (CSREES), Grant/Award Number: HAW08039-H.

## Author Contributions

Anna Halpin-McCormick - Conceptualization, Formal analysis, Writing, Revising.

Manoj Kumar Nalla - Formal analysis, Writing, Revising.

Andrew Zhang - Writing, Revising.

Nathan Fumia - Writing, Revising.

Tsung-han Lin - Writing, Revising.

Shih-wen Lin - Writing, Revising.

Yen-wei Wang - Writing, Revising.

Herbaud P.F. Zohoungbogbo - Writing, Revising.

Michael Kantar - Conceptualization, Writing, Revising.

Michael A. Gore - Conceptualization, Writing, Revising.

Derek Barchenger - Conceptualization, Writing, Revising.

## Data availability

Data and scripts are available at https://github.com/ahmccormick/Chili_Pepper_GS and Figshare https://doi.org/10.6084/m9.figshare.31435636. LLM search tool for the *Capsicum* core and global collection is available at https://github.com/GEBV-collab/GEBV_Explorer_V2.git.

## Supplemental Figures

**Figure S1.**
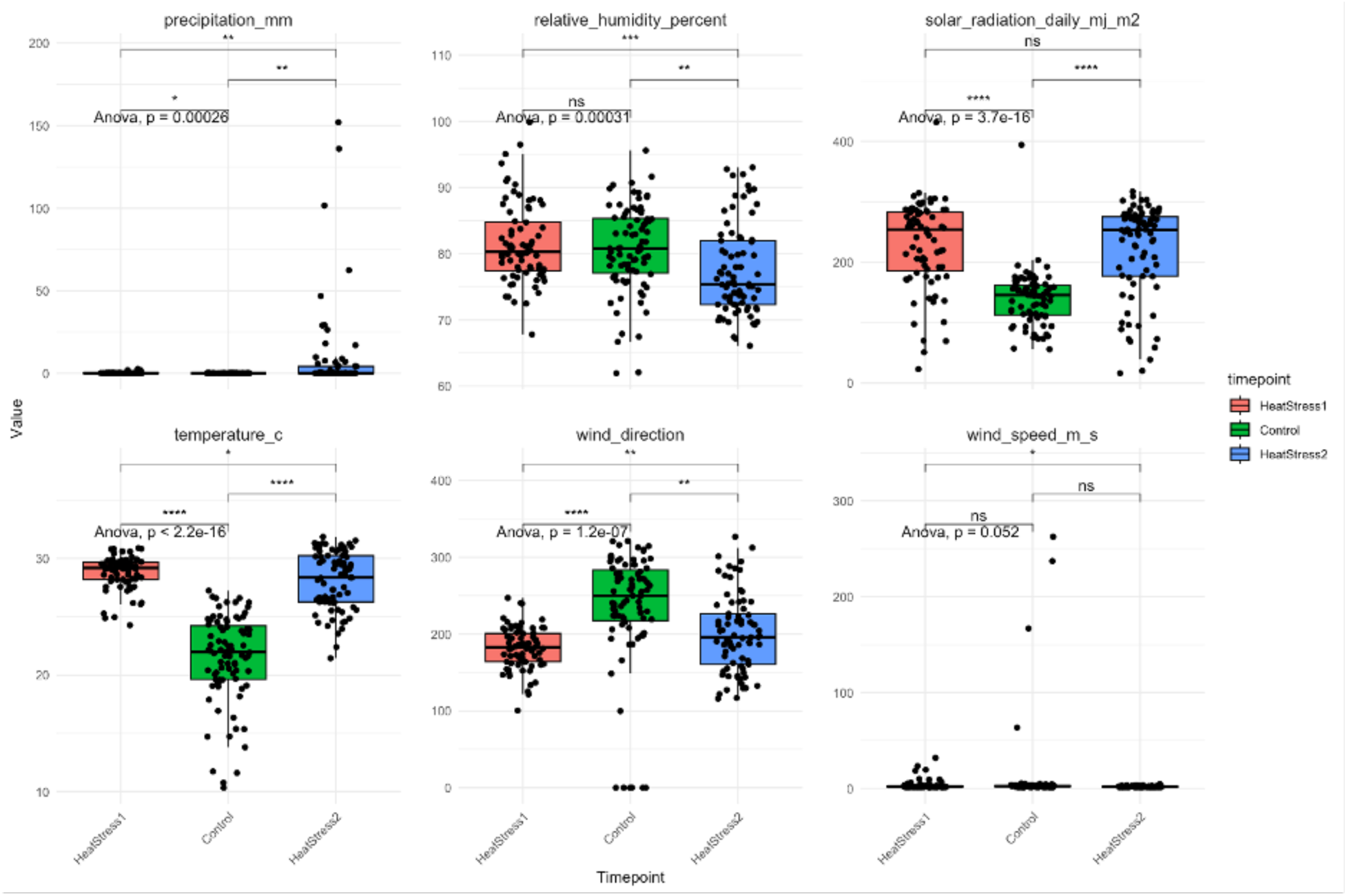
Raw climate data for the control, heat stress-1 and heat stress-2 timepoints for (**A**) precipitation (mm) (**B**) Relative humidity (%) (**C**) Solar radiation (mj/m^2^) (**D**) Temperature (°C) (**E**) Wind direction (**F**) Wind speed (m/s)

**Figure S2.**
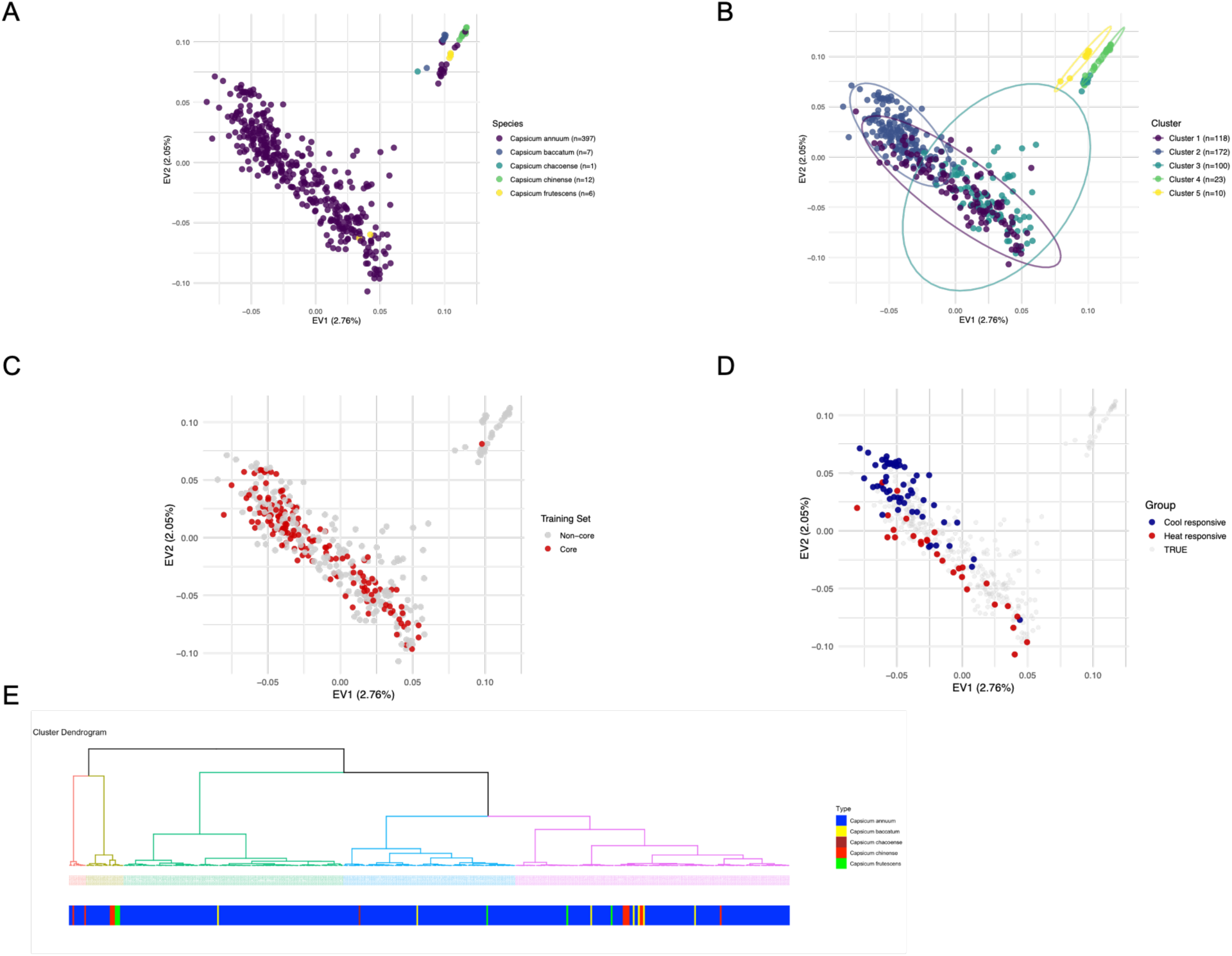
Population structure for the core (n=423) *Capsicum* collection (**A**) colored by species **(B**) coloured by the 5 groups identified by hierarchical clustering of principal components (HCPC) (**C**) coloured by training samples (n=150, red) and test samples (grey) (**D**) heat responsive (n=28, red) and cool responsive (n=57, blue) (**E**) HCPC reveals 5 main groups, coloured by species.

**Figure S3.**
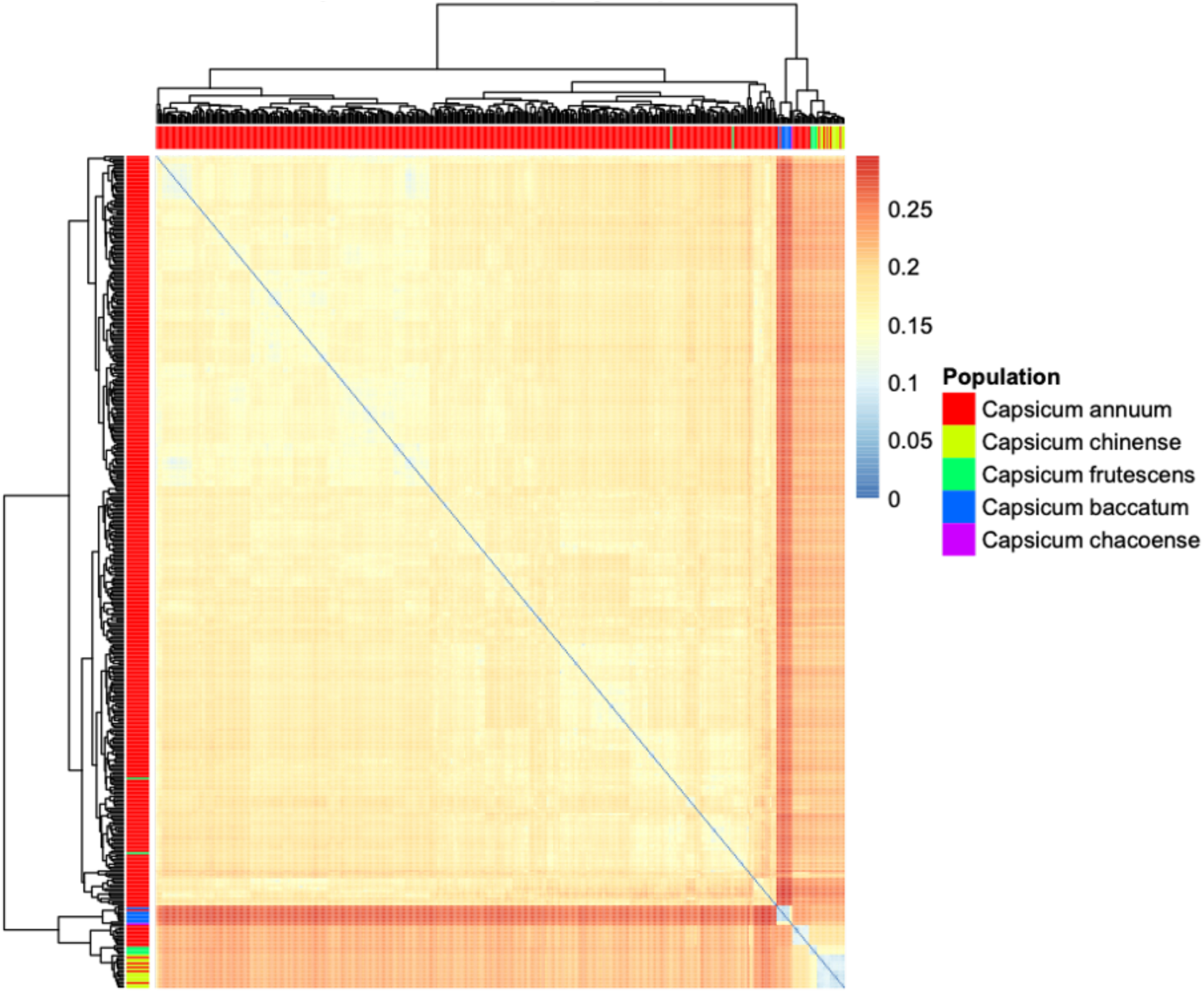
Kinship matrix heatmap for the core collection (n=423) coloured by species.

**Figure S4.**
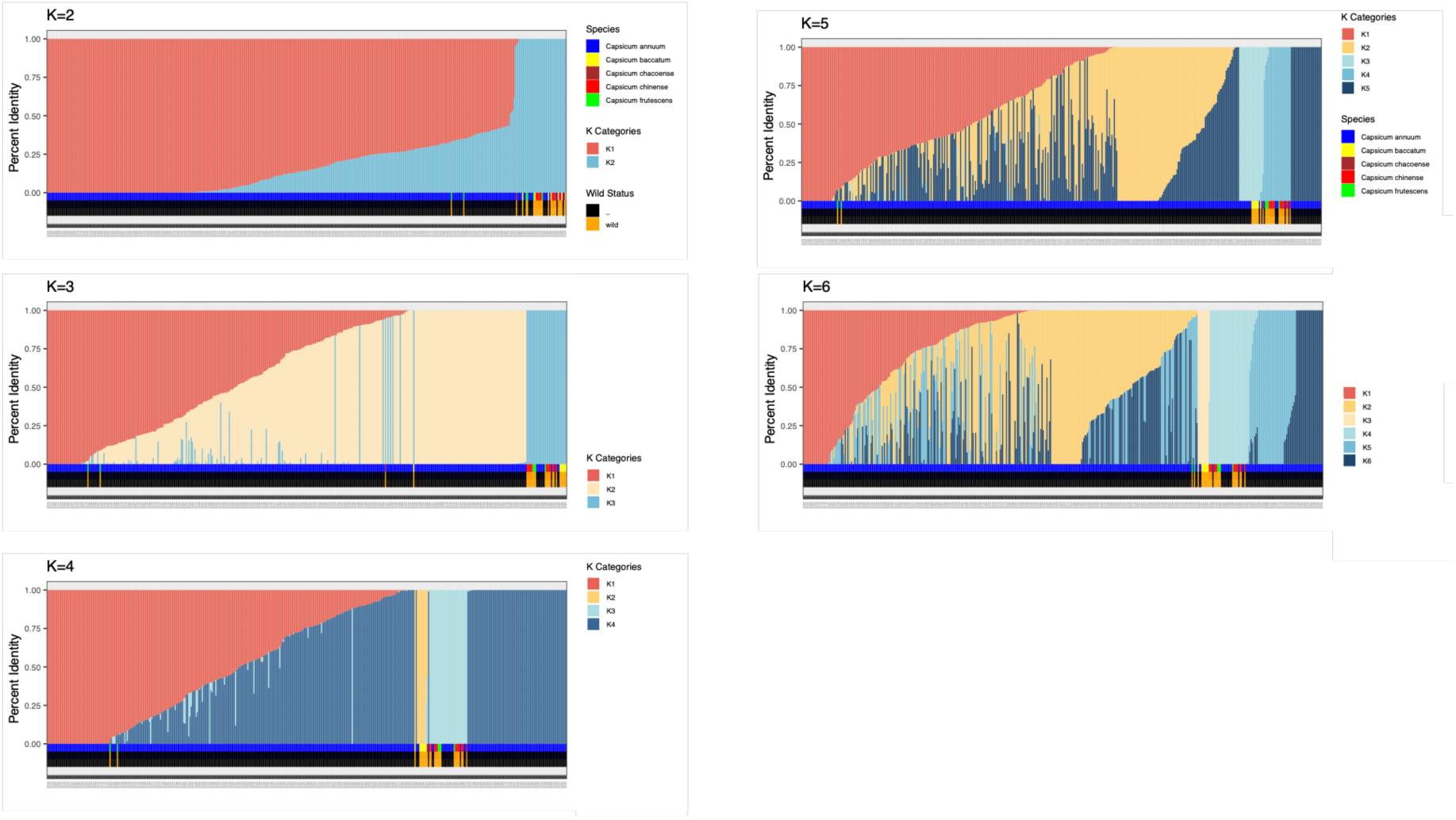
fastSTRUCTURE (K2-6) for the core collection (n=423) using 340,734 SNPs, coloured by species.

**Figure S5.**
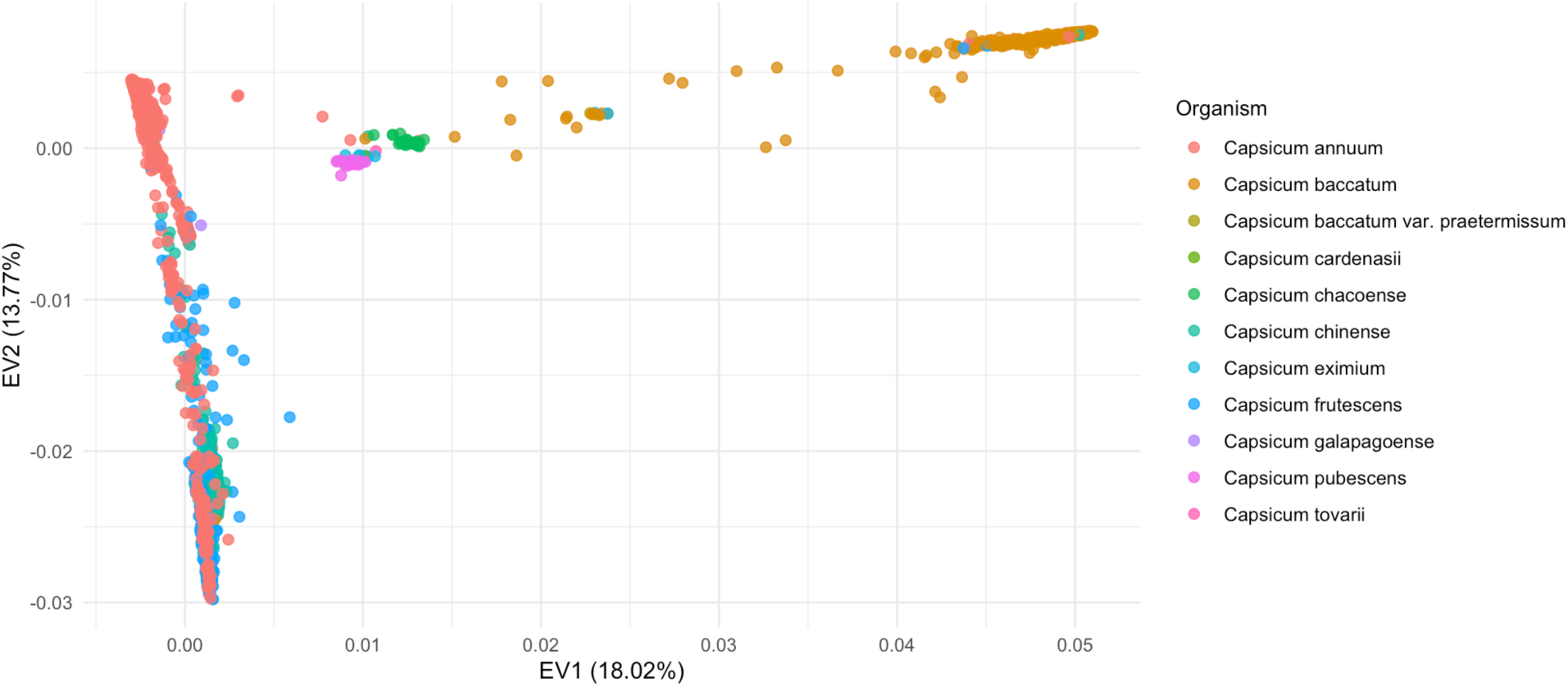
PCA for the global *Capsicum* collection (n=10,250) coloured by species.

**Figure S6.**
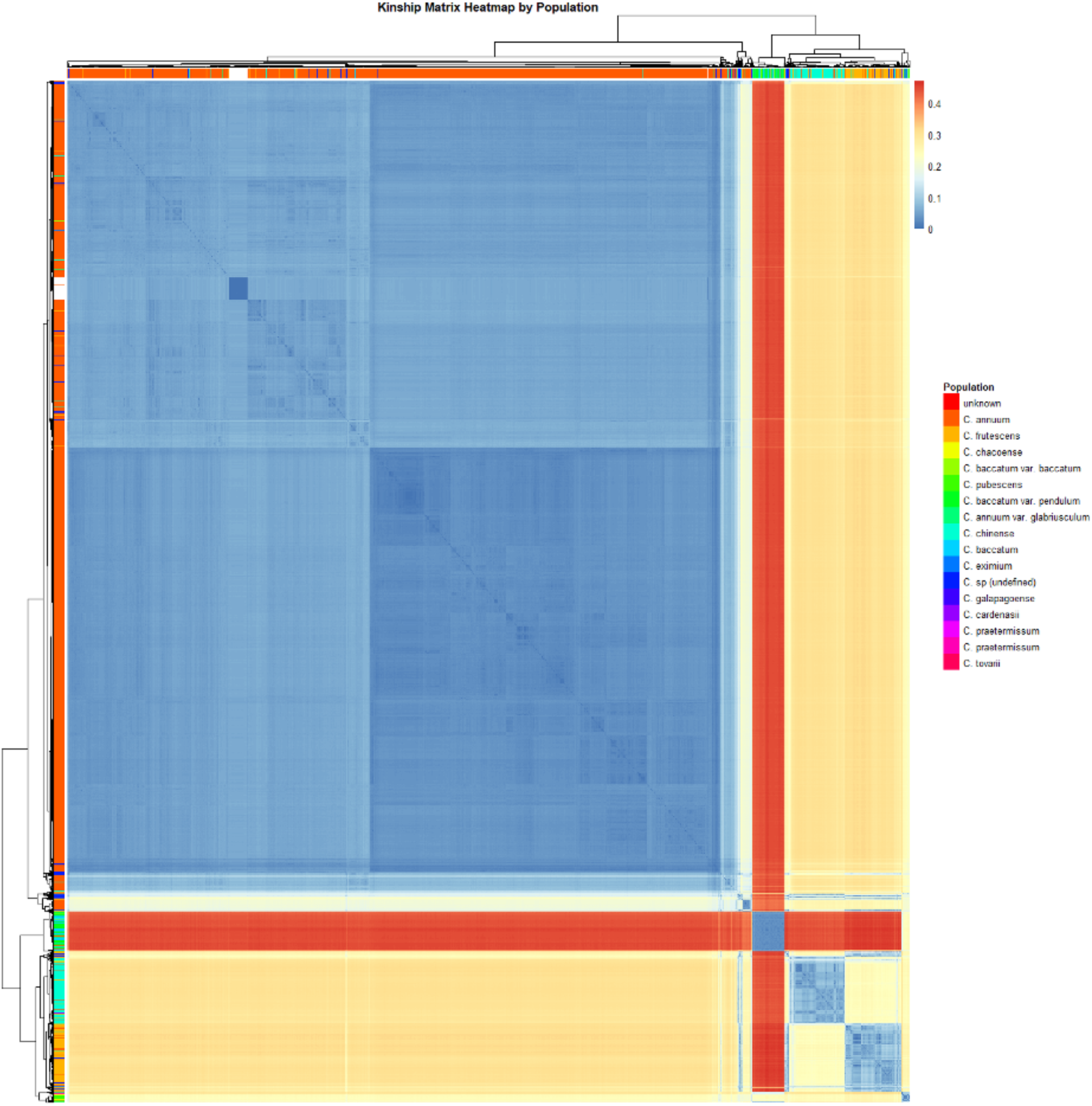
Kinship matrix heatmap for the global collection (n=10,250) coloured by species.

**Figure S7.**
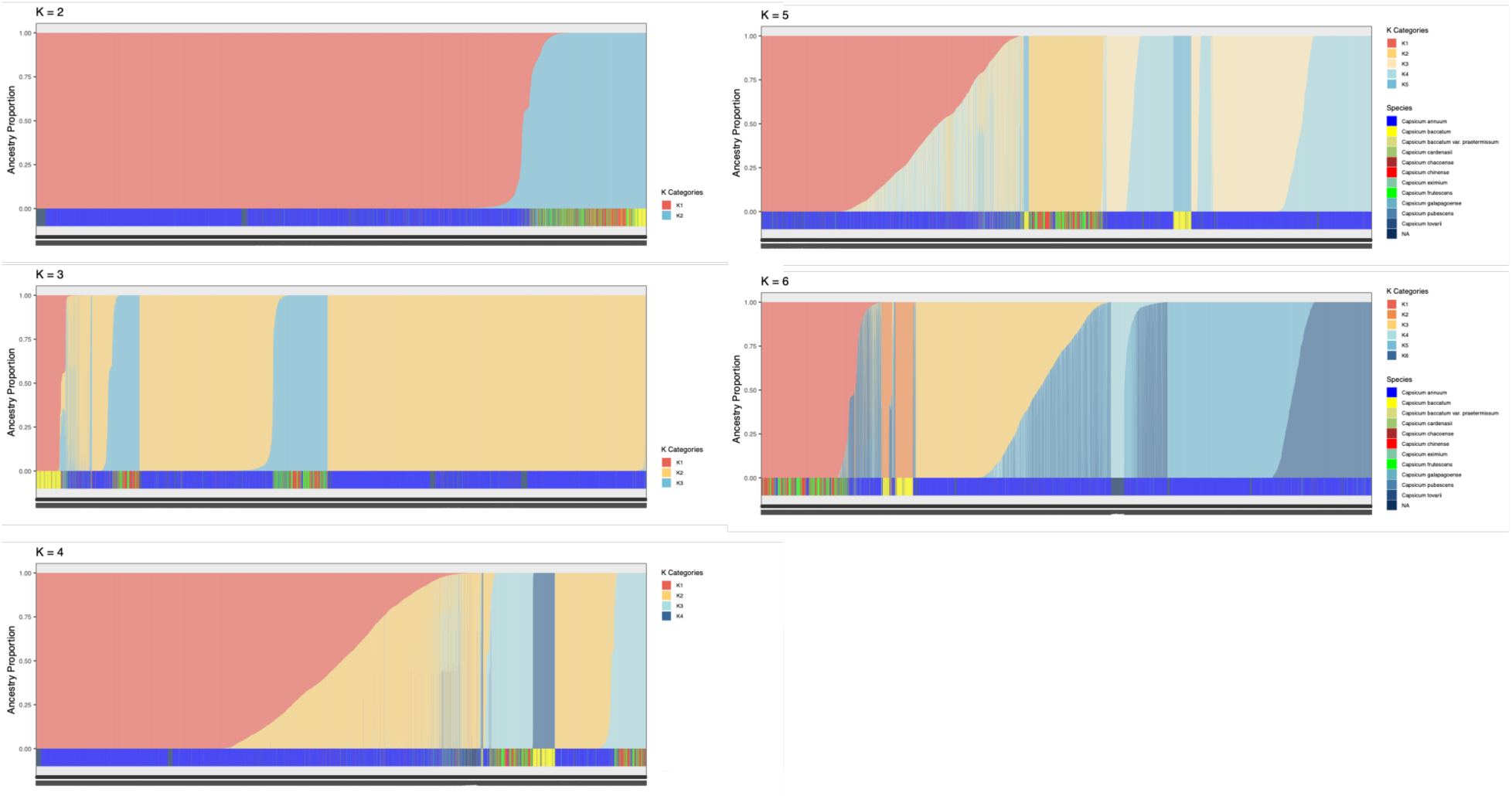
fastSTRUCTURE for global collection (n=10,250) using 23,462 SNPs and coloured by species.

**Figure S8.**
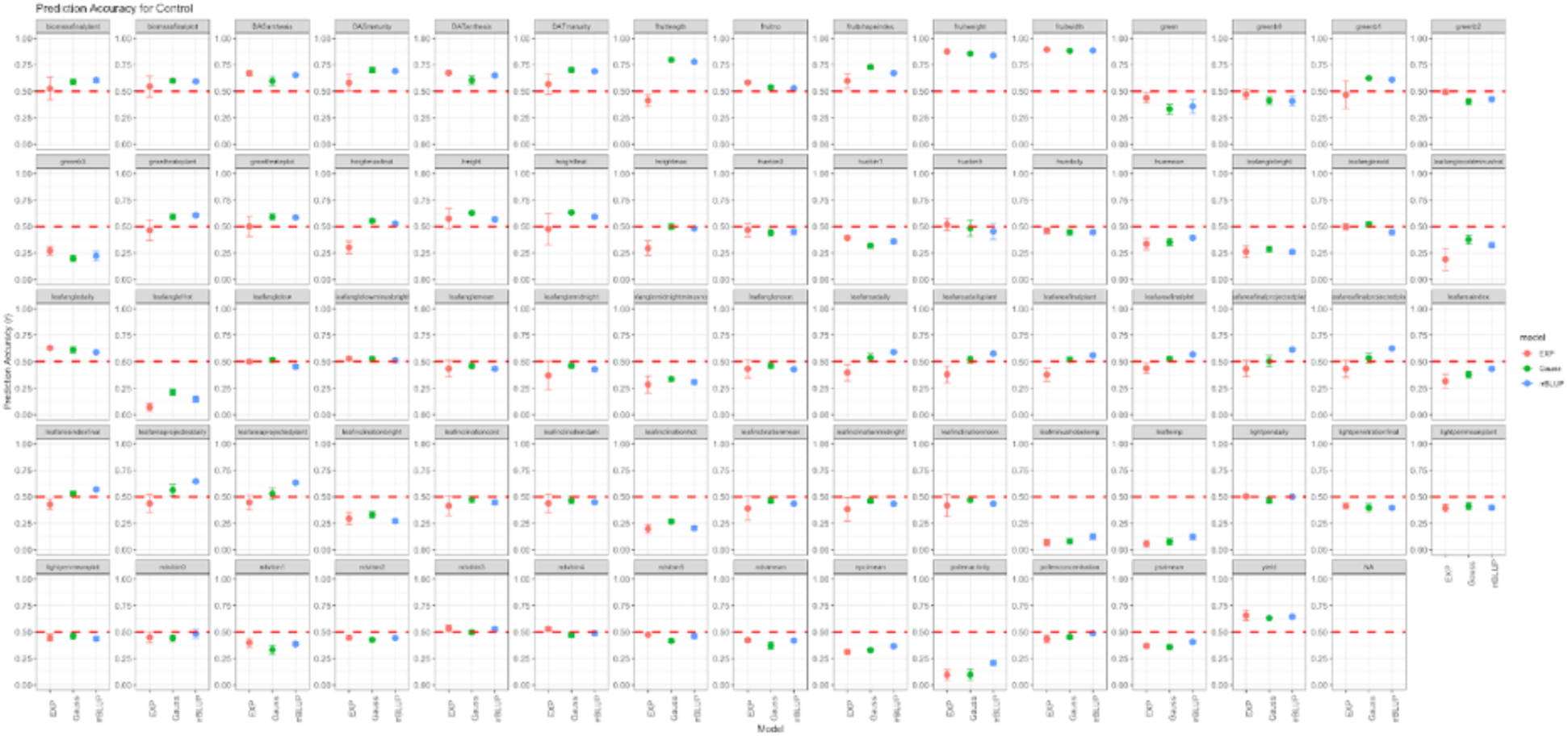
73 trait prediction accuracies for control timepoint for the core collection.

**Figure S9.**
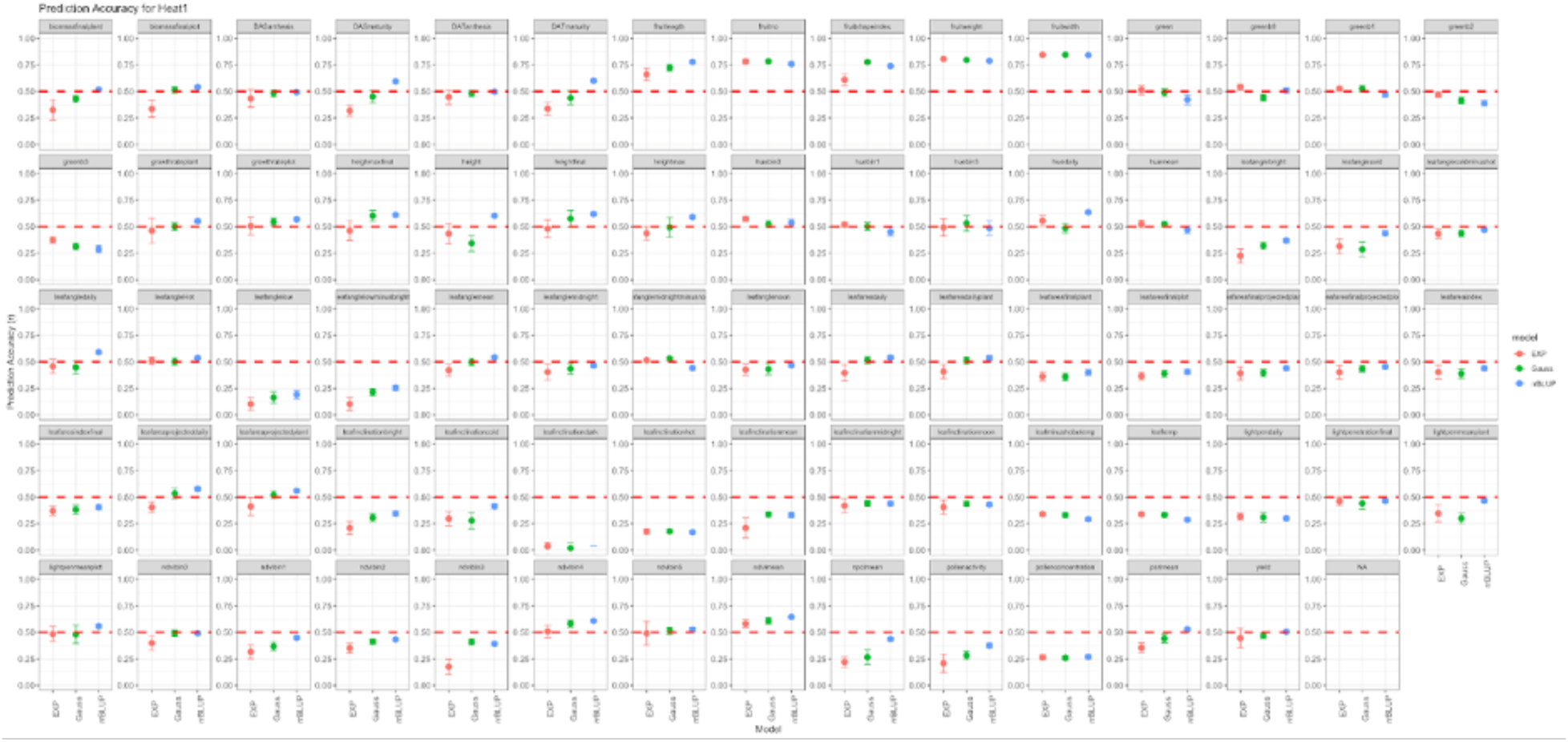
73 trait prediction accuracies for heat stress #1 timepoint for the core collection.

**Figure S10.**
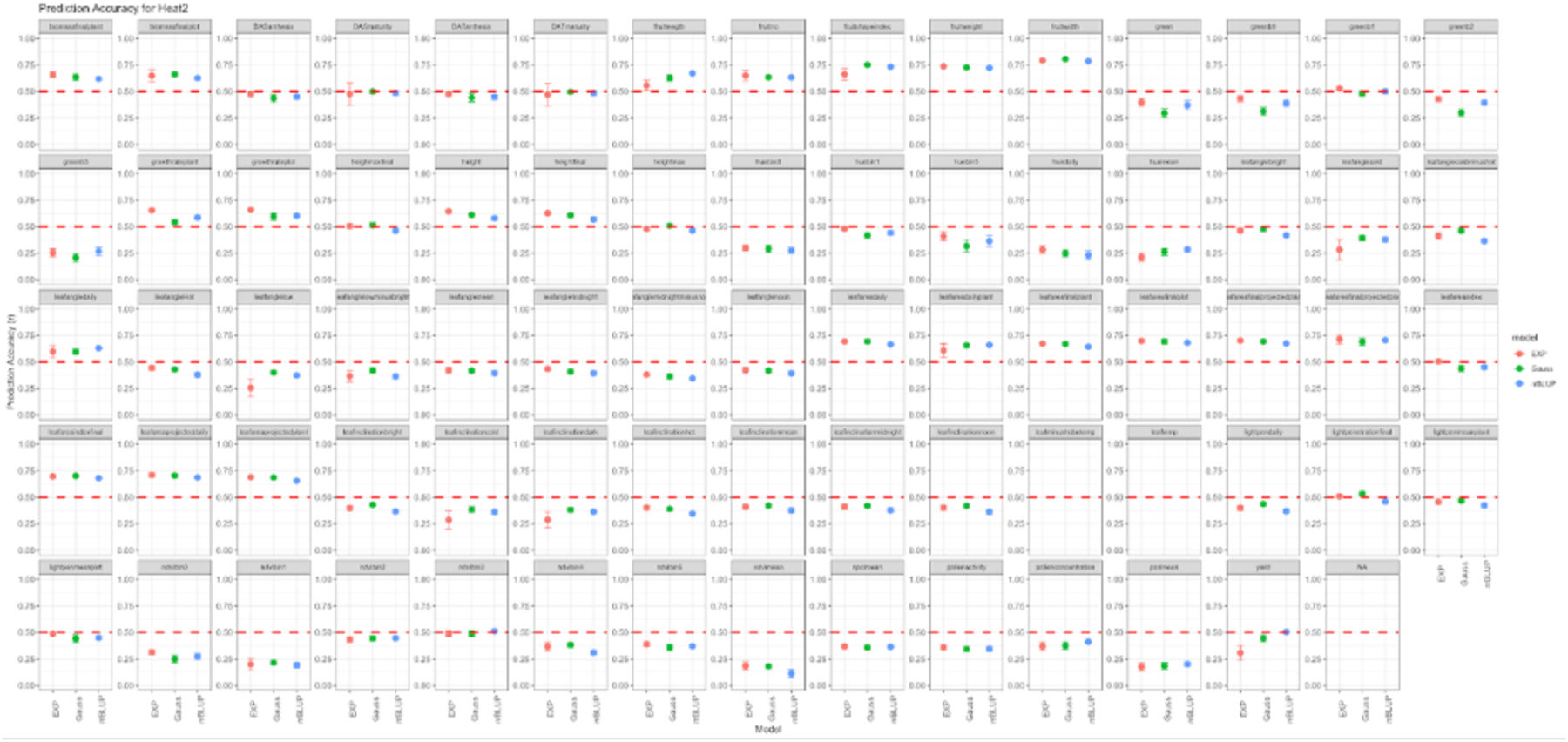
73 trait prediction accuracies for heat stress #2 timepoint for the core collection.

**Figure S11.**
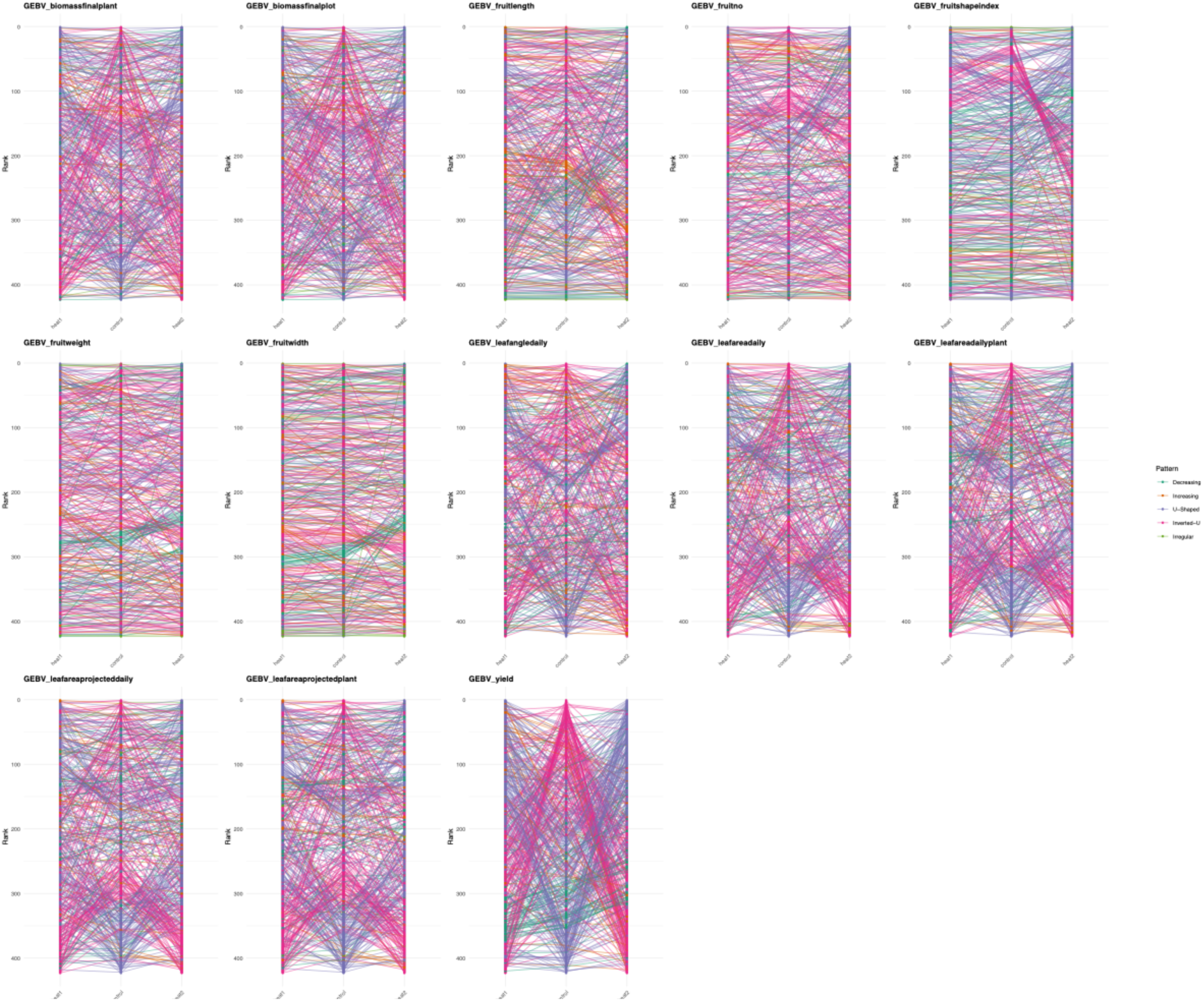
Line rank changes for traits (n=13) with prediction accuracies above 0.5 for rrBLUP across all three environmental conditions for the core collection.

**Figure S12.**
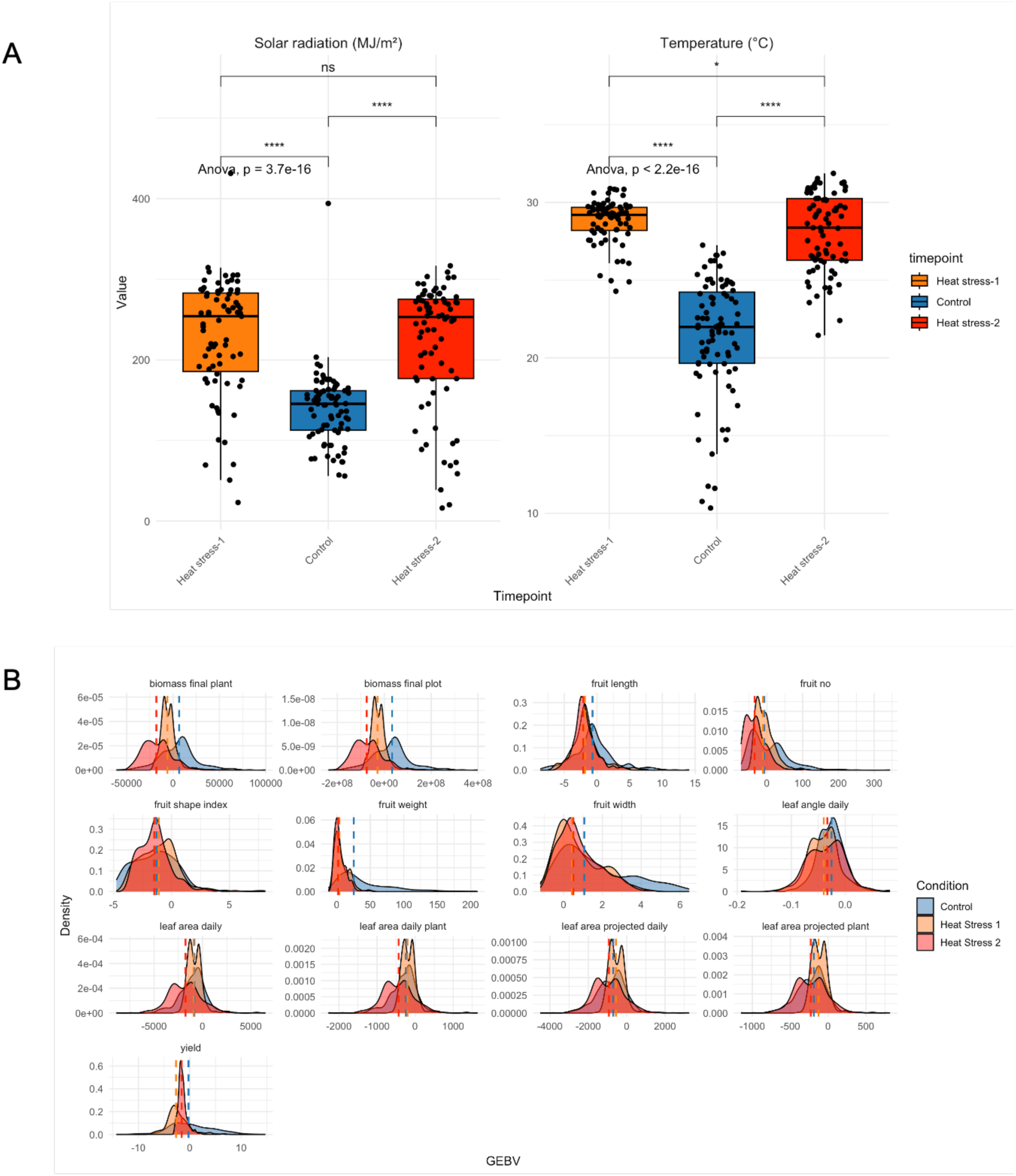
Climate differences in experimental timepoints affect GEBV distributions (**A**) Raw climate data for the control, heat stress-1 and heat stress-2 timepoints for Solar radiation (mj/m^2^) and Temperature (°C) (**B**) Shifts in agronomic trait GEBV distributions in core collection for 13 traits with prediction accuracies above 0.5 across all three conditions. The dashed line is the median line for the GEBV for a timepoint.

**Figure S13.**
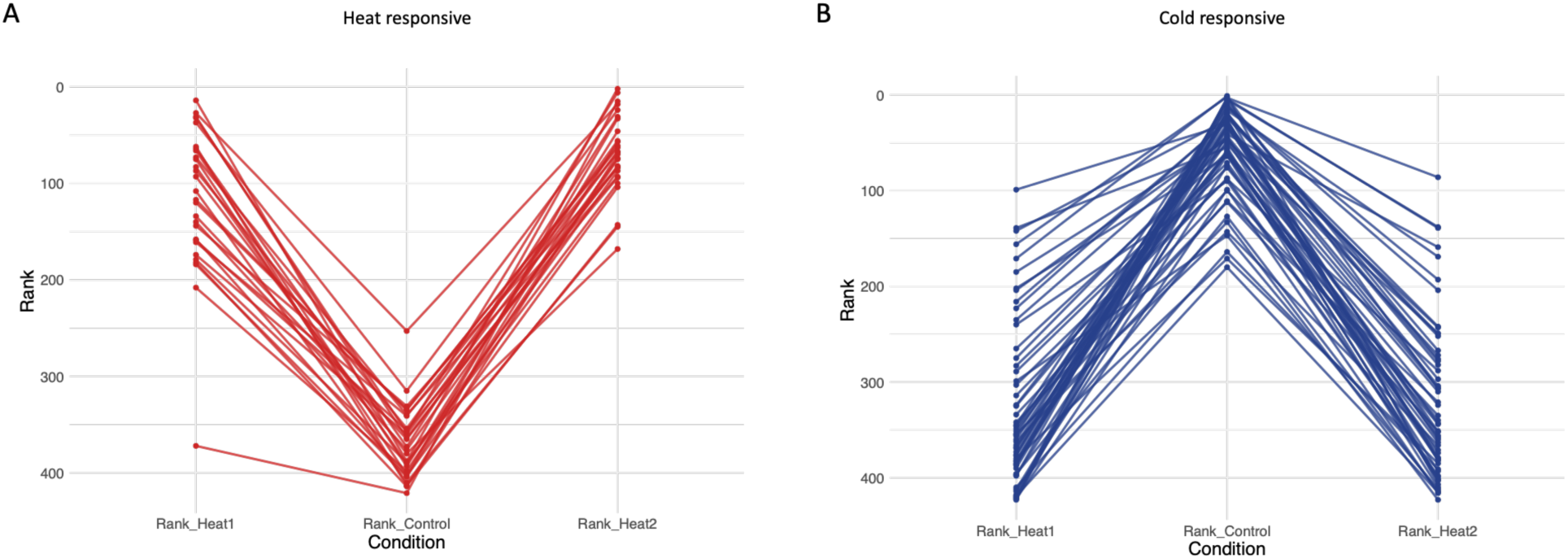
Evidence of heat responsiveness in the *Capsicum* core collection using Genomic Selection (GS). Genotype rank changes between heat stress-1, control and heat stress-2 for fruit yield (PA = 0.56, 0.61, 0.54 respectively). Rank changes for GEBVs in response to heat stress across 423 *Capsicum* accessions for fruit yield (tonnes/ha). Accessions which show largest rank changes (top 20%) in yield with respect to heat stress-1, control and heat stress-2 conditions were selected (**A**) Potential heat-tolerant accessions (n = 28/203) (**B**) Potential accessions (n = 57/155) with a preference for colder control temperatures.

**Figure S14.**
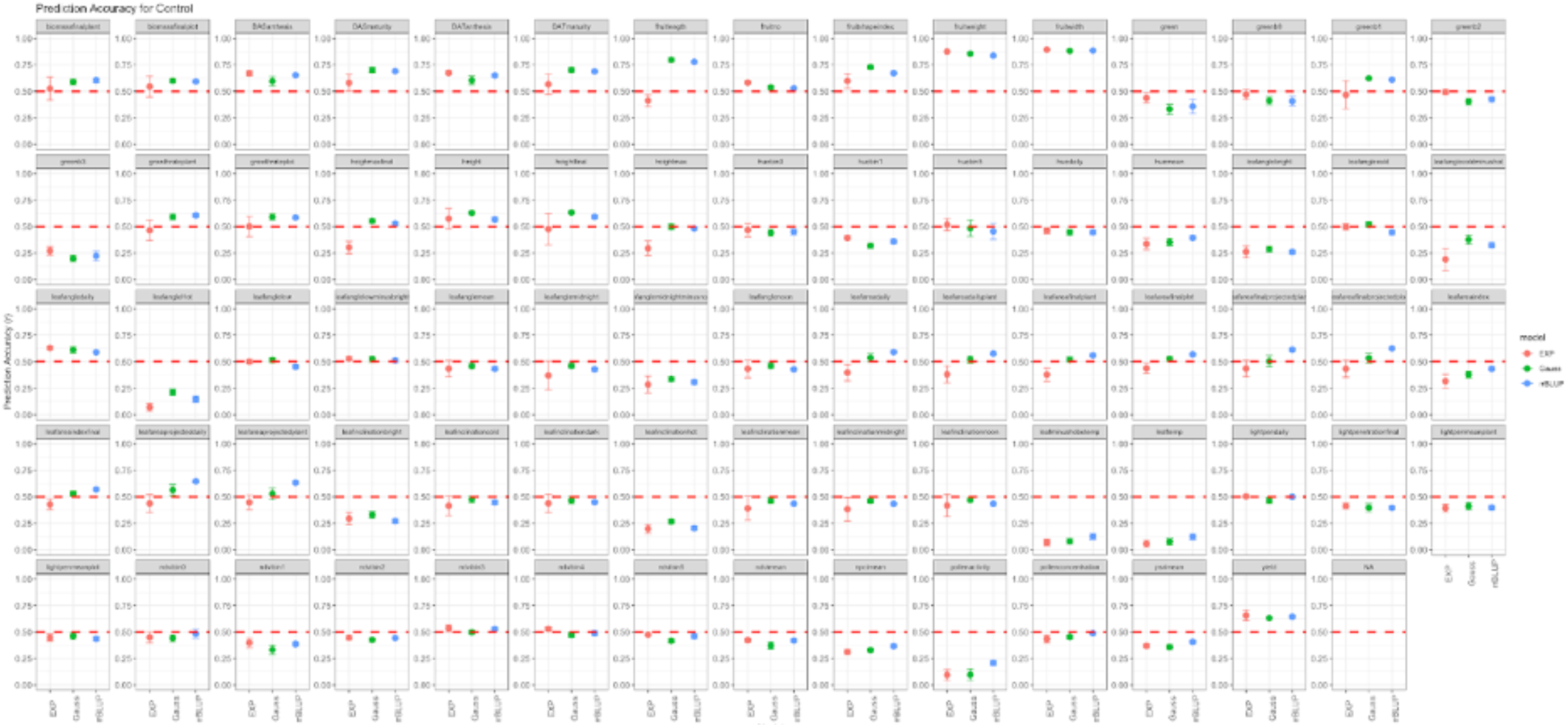
73 trait prediction accuracies for control timepoint in the global collection.

**Figure S15.**
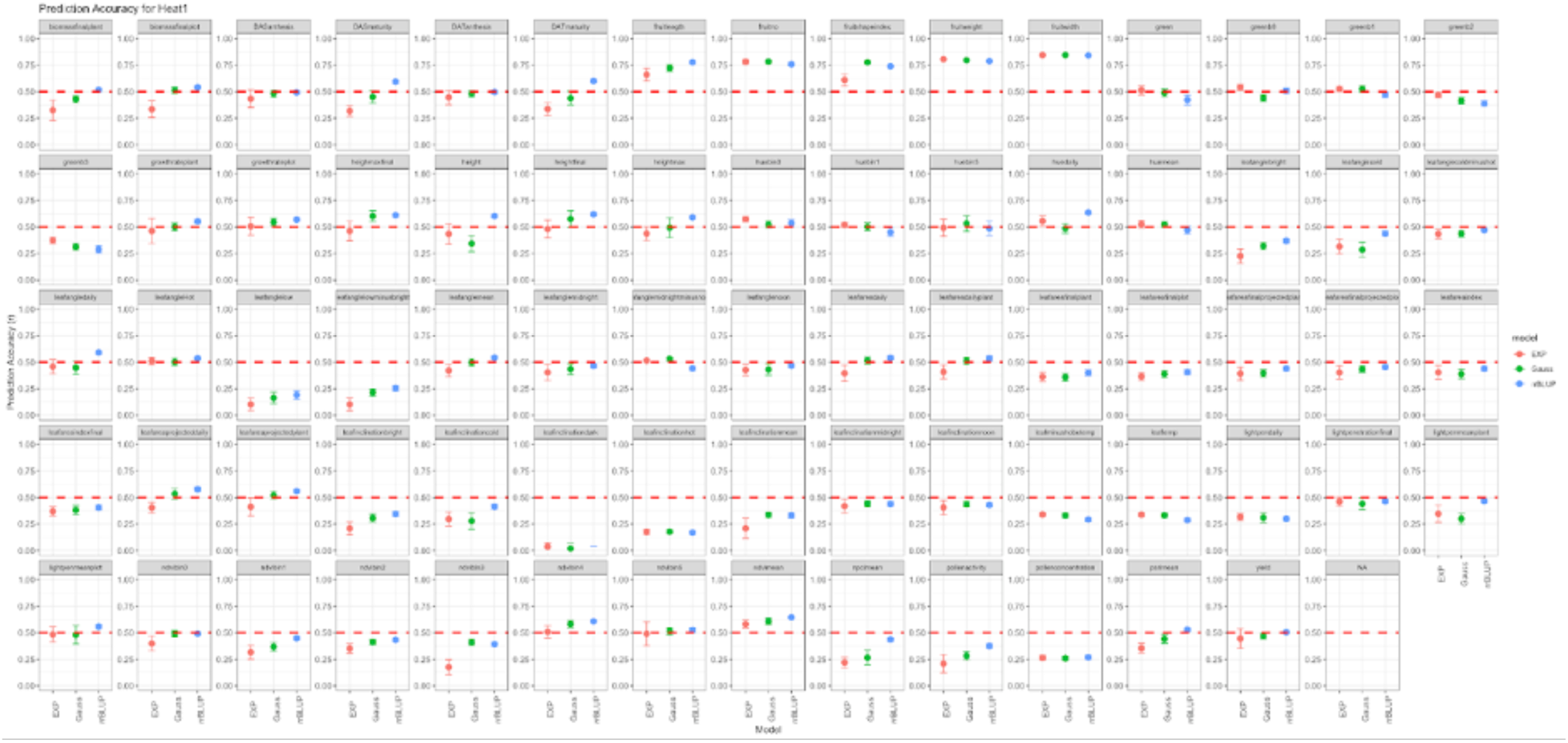
73 trait prediction accuracies for heat stress #1 timepoint in the global collection.

**Figure S16.**
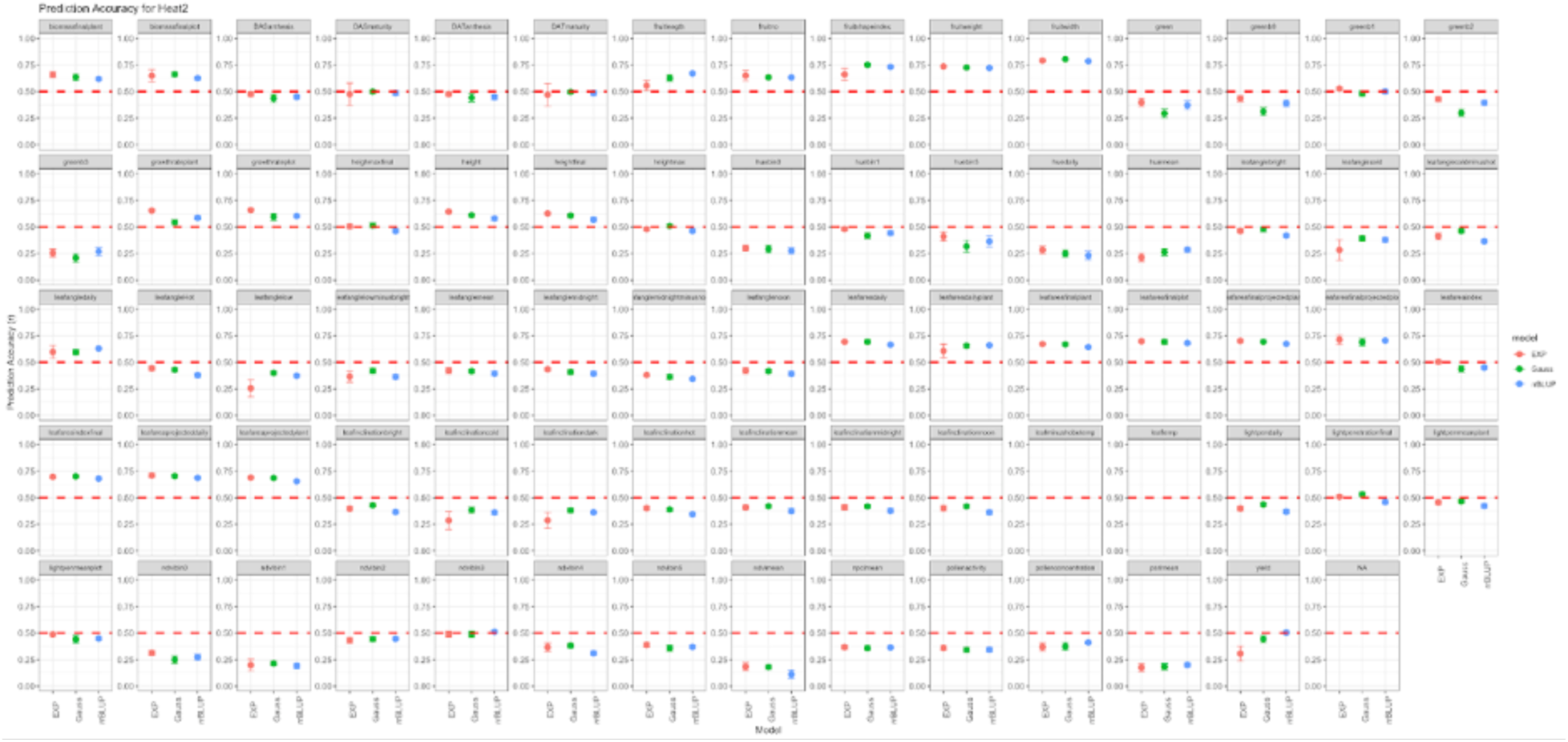
73 trait prediction accuracies for heat stress #2 timepoint in the global collection.

**Figure S17.**
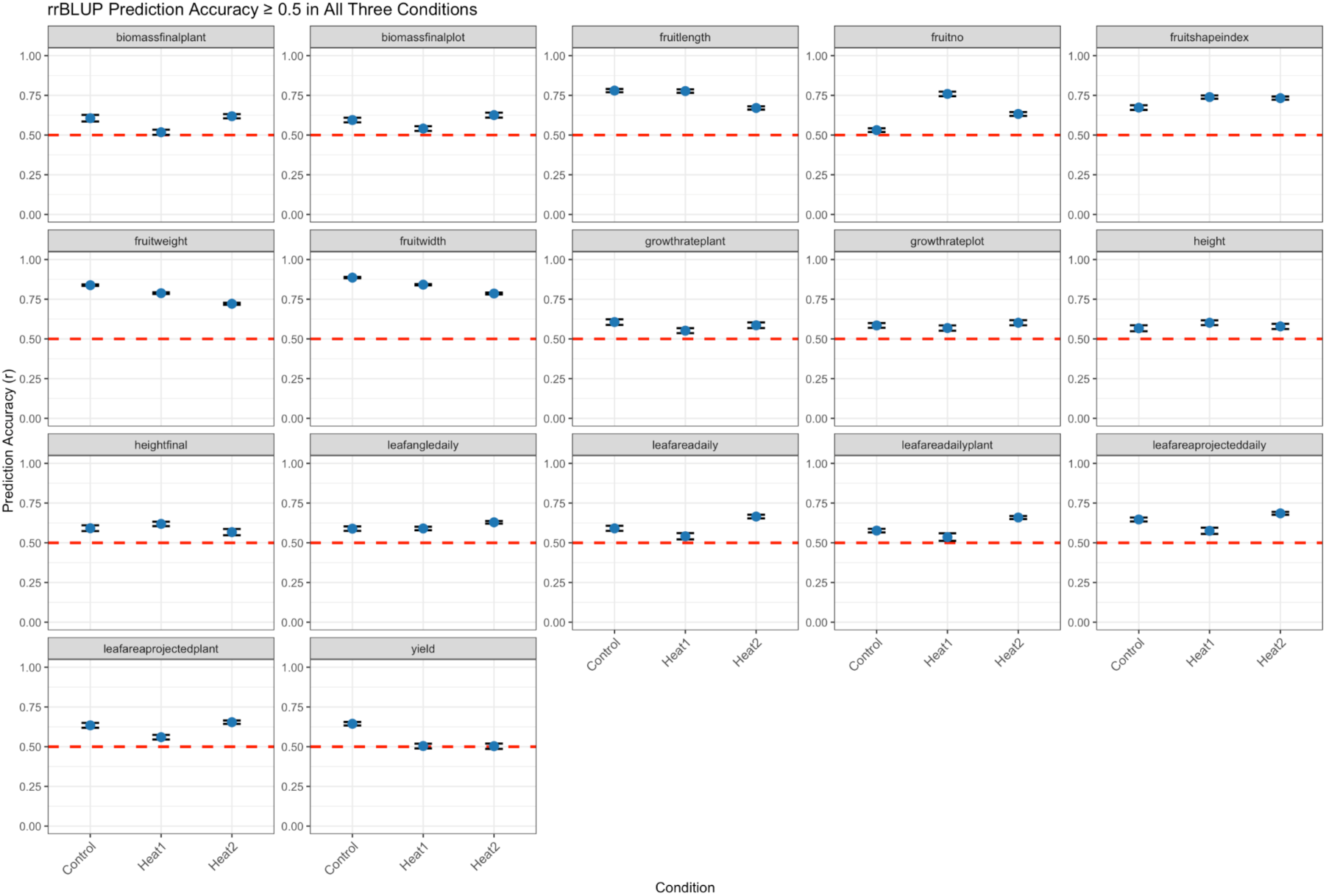
Global results for the 17 traits with prediction accuracies above 0.5 for rrBLUP.

**Figure S18.**
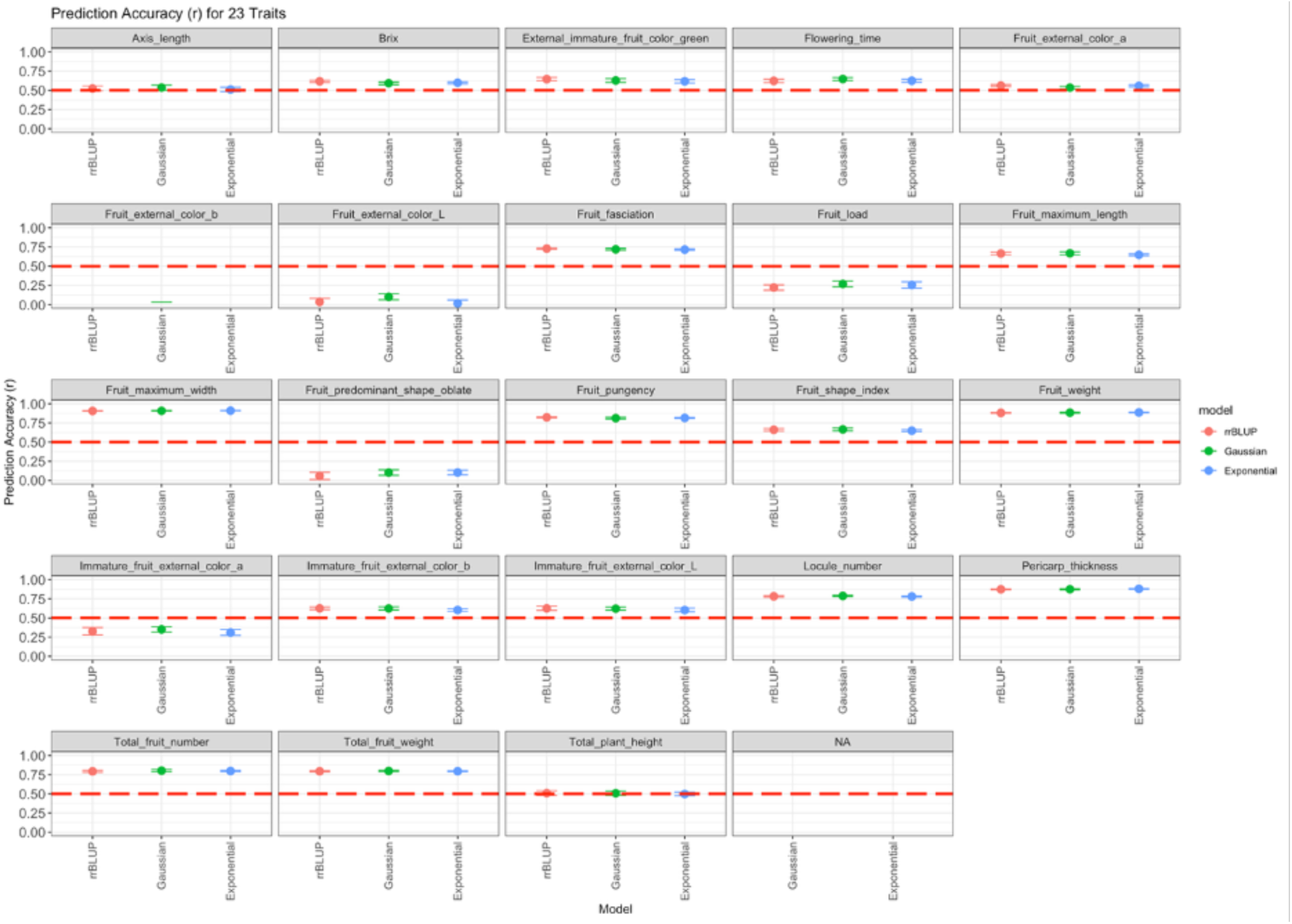
23 quality trait prediction accuracies for chili core collection (18 traits over 0.5).

**Figure S19.**
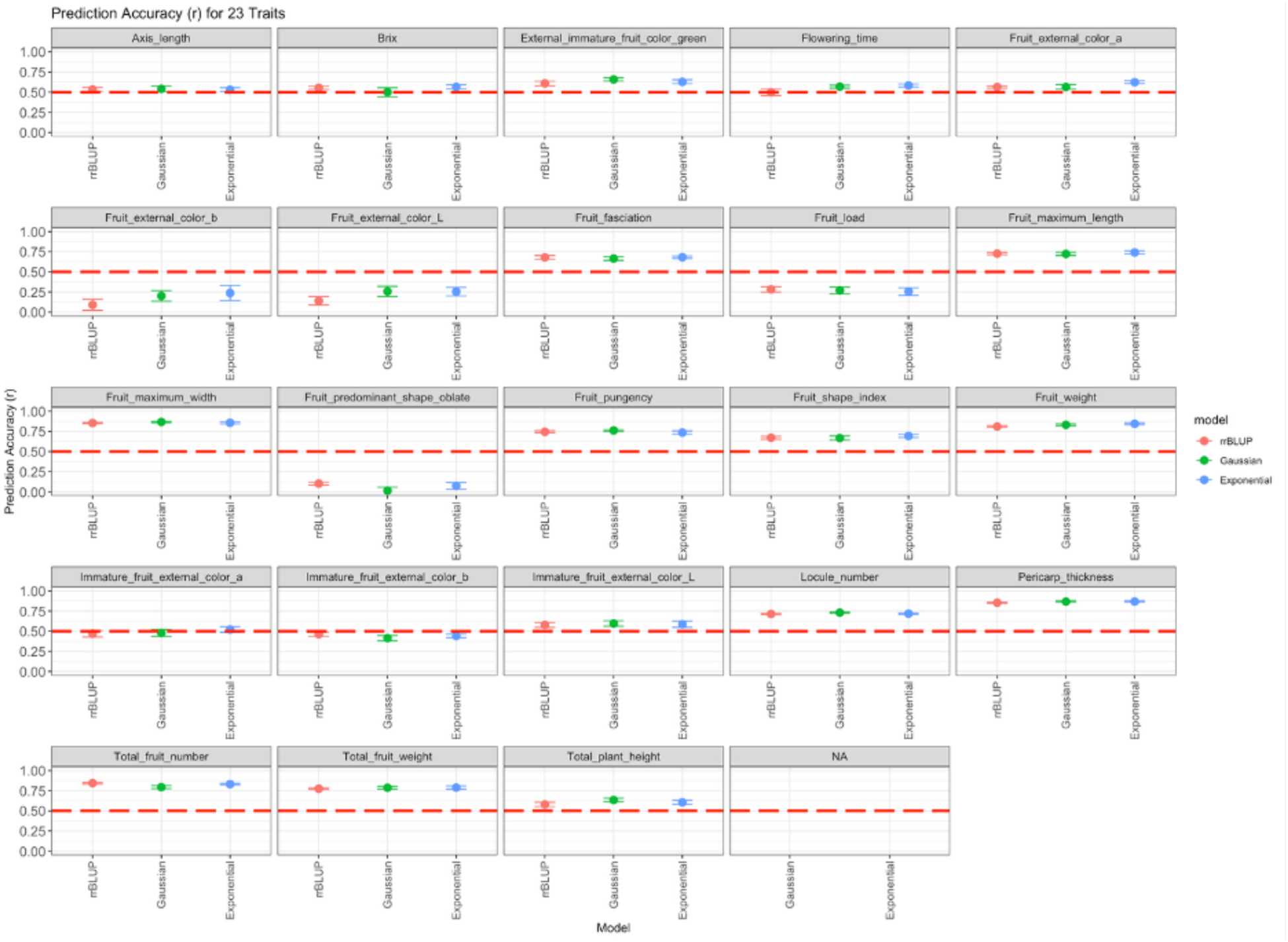
23 quality trait prediction accuracies for chili global 10k collection (17 traits over 0.5).

**Figure S20.**
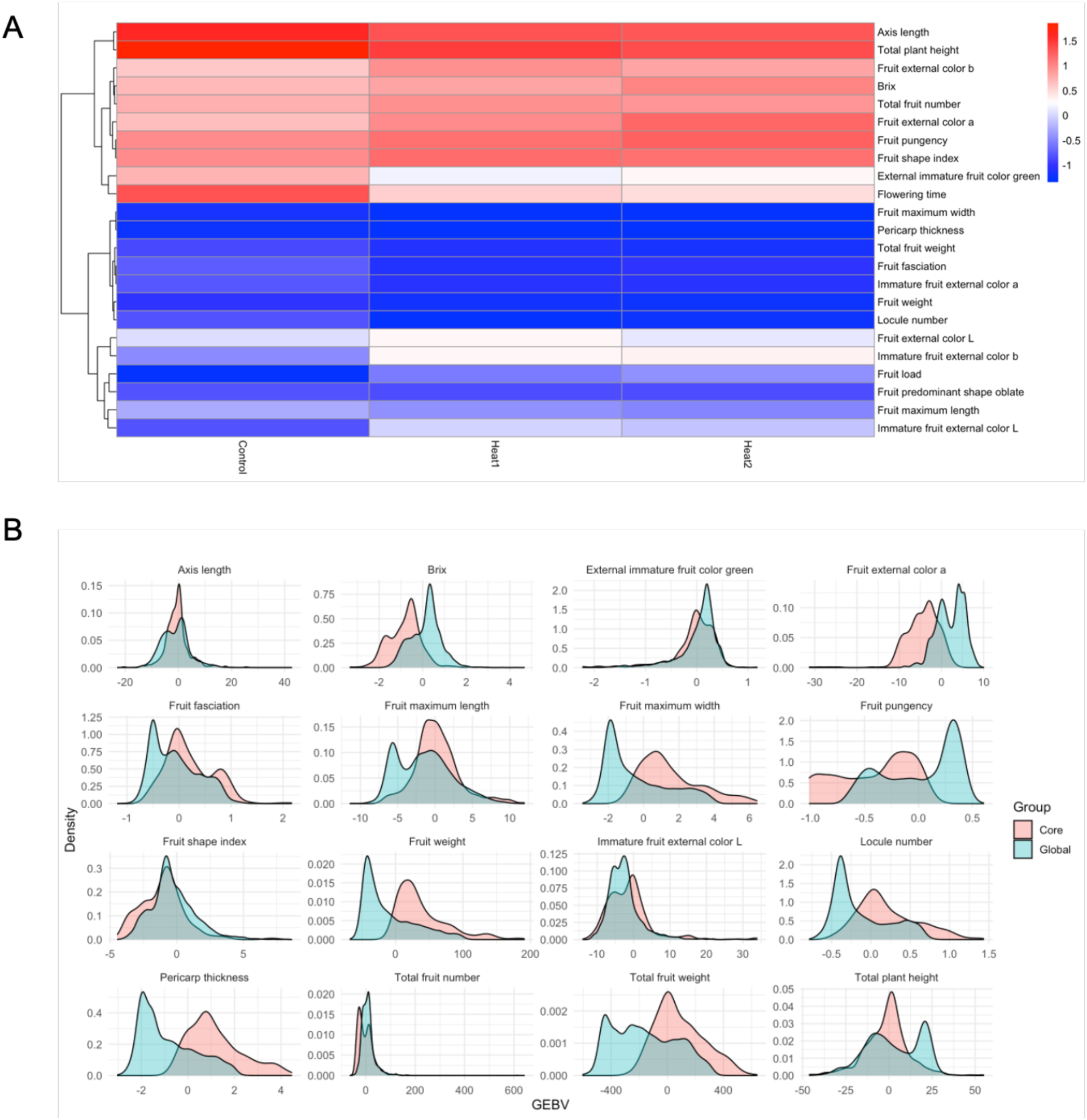
Relationship between climate and quality trait GEBV distributions in core and global *Capsicum* collections (**A**) Climate and quality trait relationships for environment in the core collection (n=423). All 73 climate traits averaged for a per line GEBV average representing the environment (control, heat stress-1 or 2). Positively correlated breeding values for a subset of quality traits and environment observed (**B**) GEBV distributions for the 16 quality traits with prediction accuracies above 0.5 in core and global collections.

**Figure S21.**
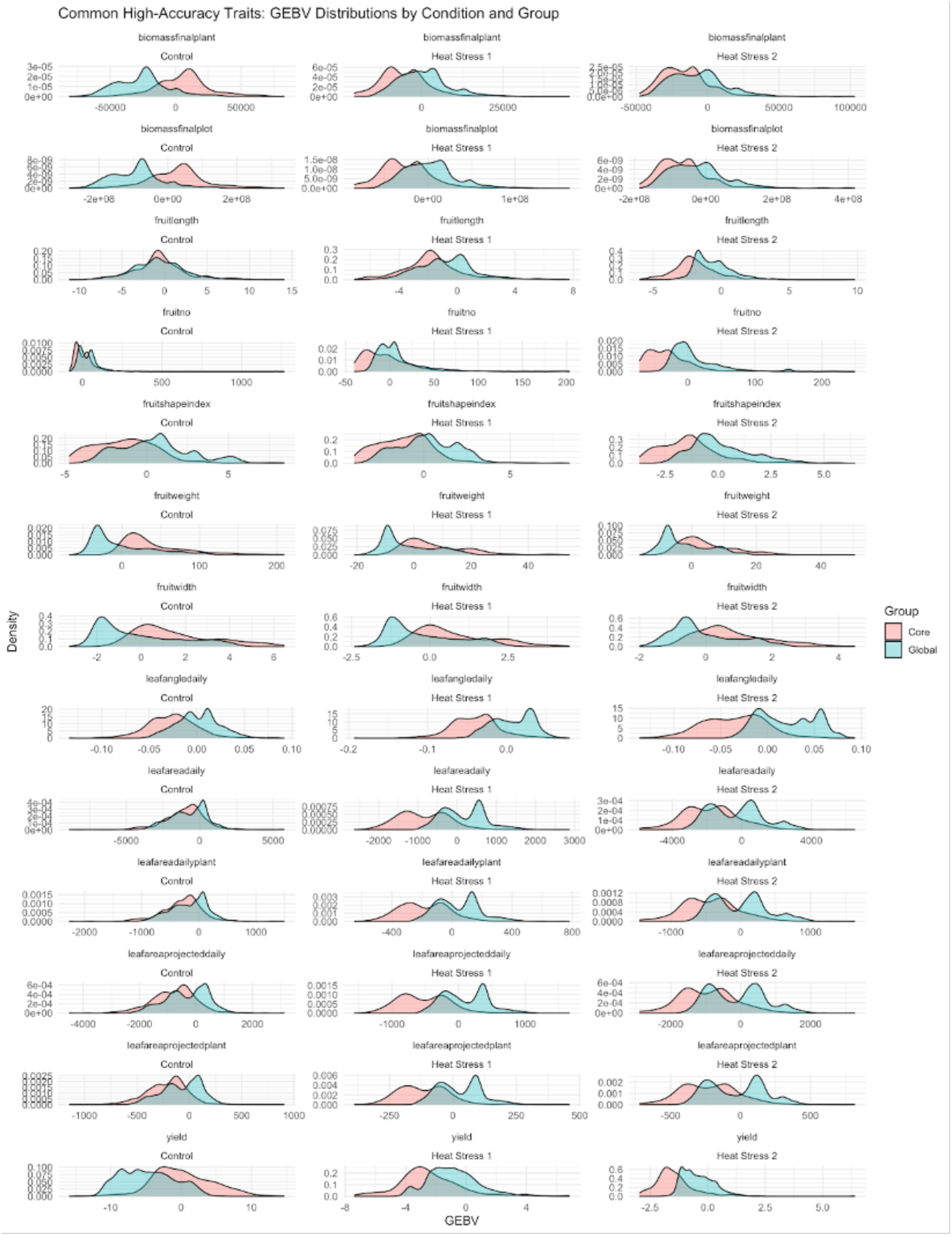
Agronomic trait GEBV distributions in core and global collections. There were 13 traits with prediction accuracies above 0.5 in core and global collections across all three conditions.

**Figure S22.**
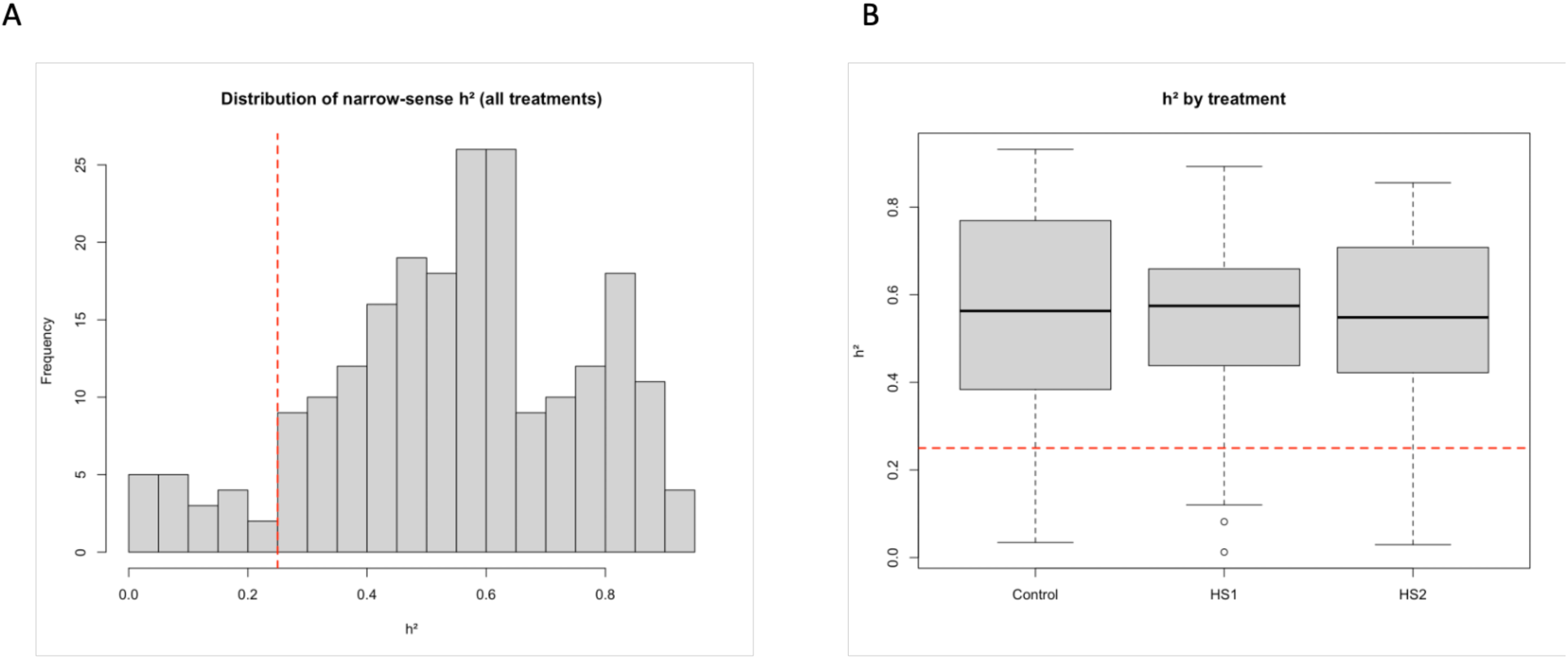
Understanding the relationship between heritability and prediction accuracy. Distribution of genomic narrow sense heritability (h^2^) across traits and conditions (**A**) Histogram of h^2^ estimates for all traits across control, heat stress-1 and heat stress-2 (**B**) Boxplot of h^2^ estimates by condition.

**Figure S23.**
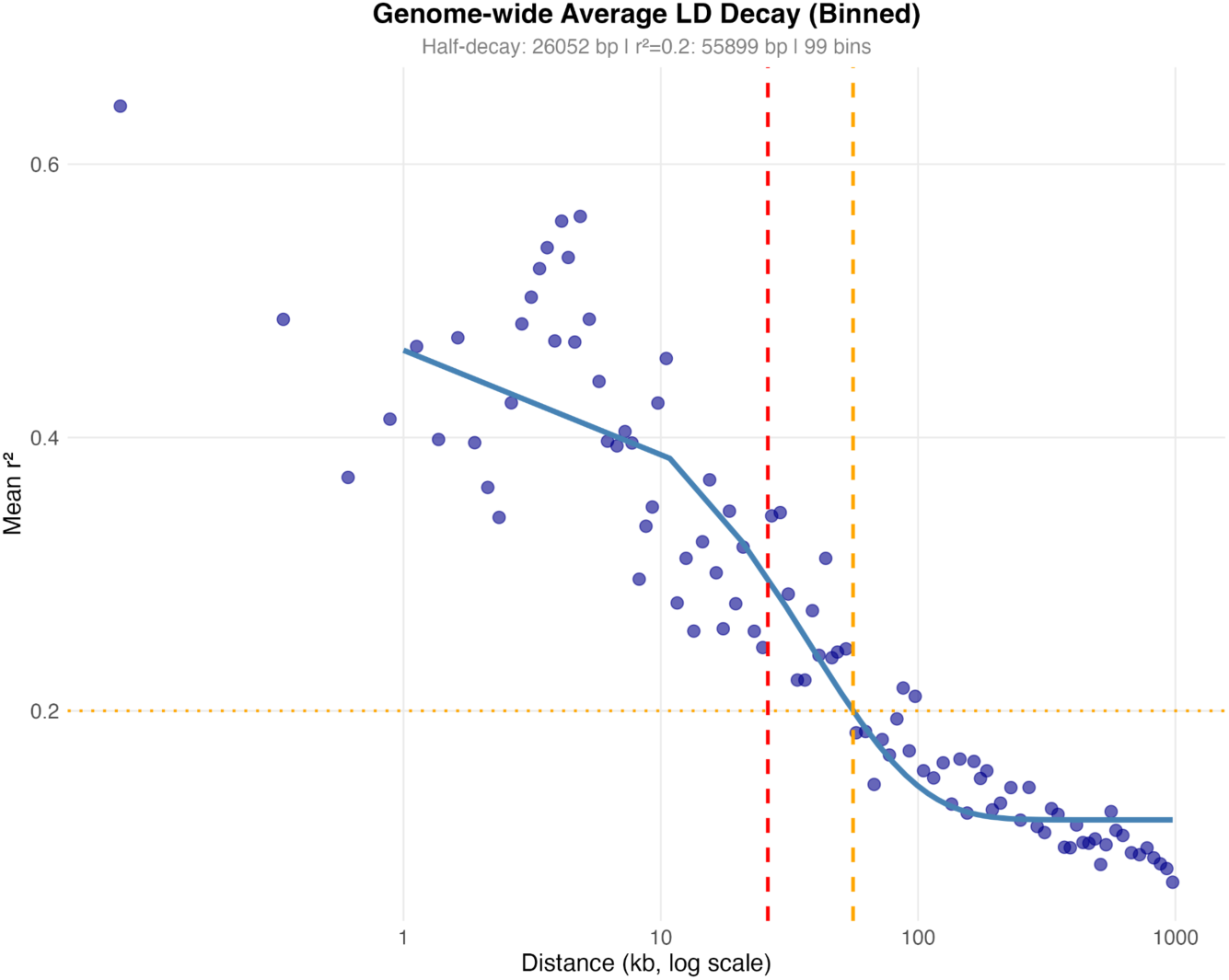
LD decay across the genome in the core collection. Here we see that LD decays to r = 0.2 at 56 kbp.

**Figure S24.**
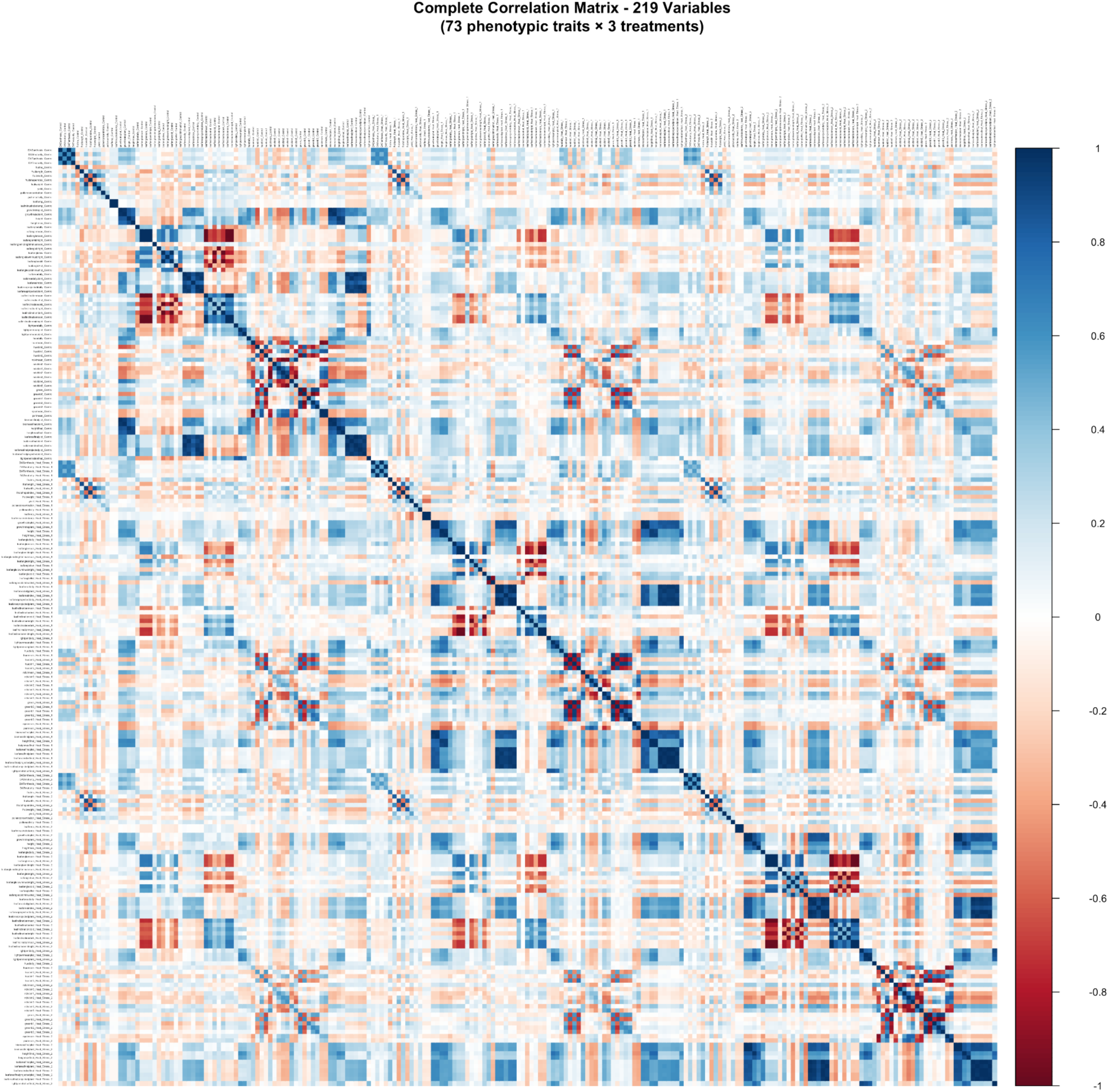
Correlation matrix across all phenotypes and environments within the core collection of *Capsicum*.

**Figure S25.**
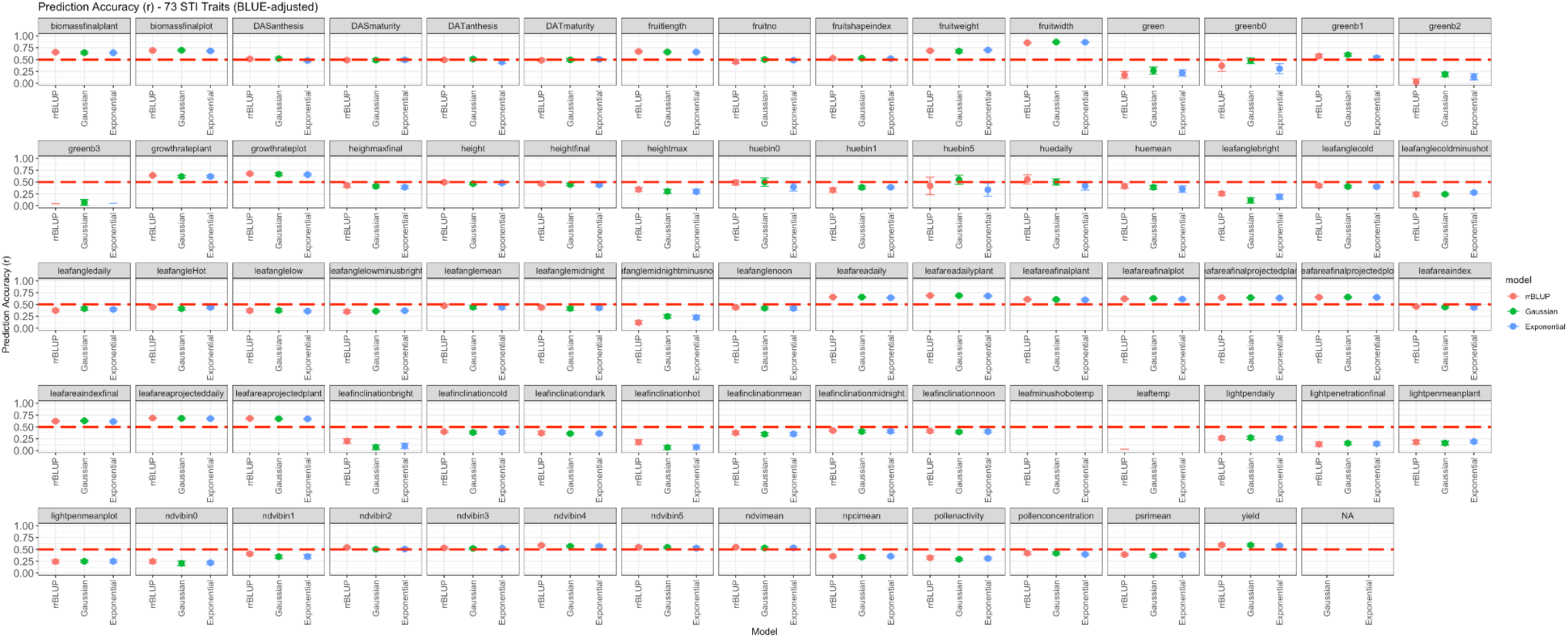
Prediction Accuracies for GS on STI index for 73 agronomic traits for the *Capsicum* core collection. 24 traits were above 0.5 prediction accuracy across all 3 models tested.

**Figure S26.**
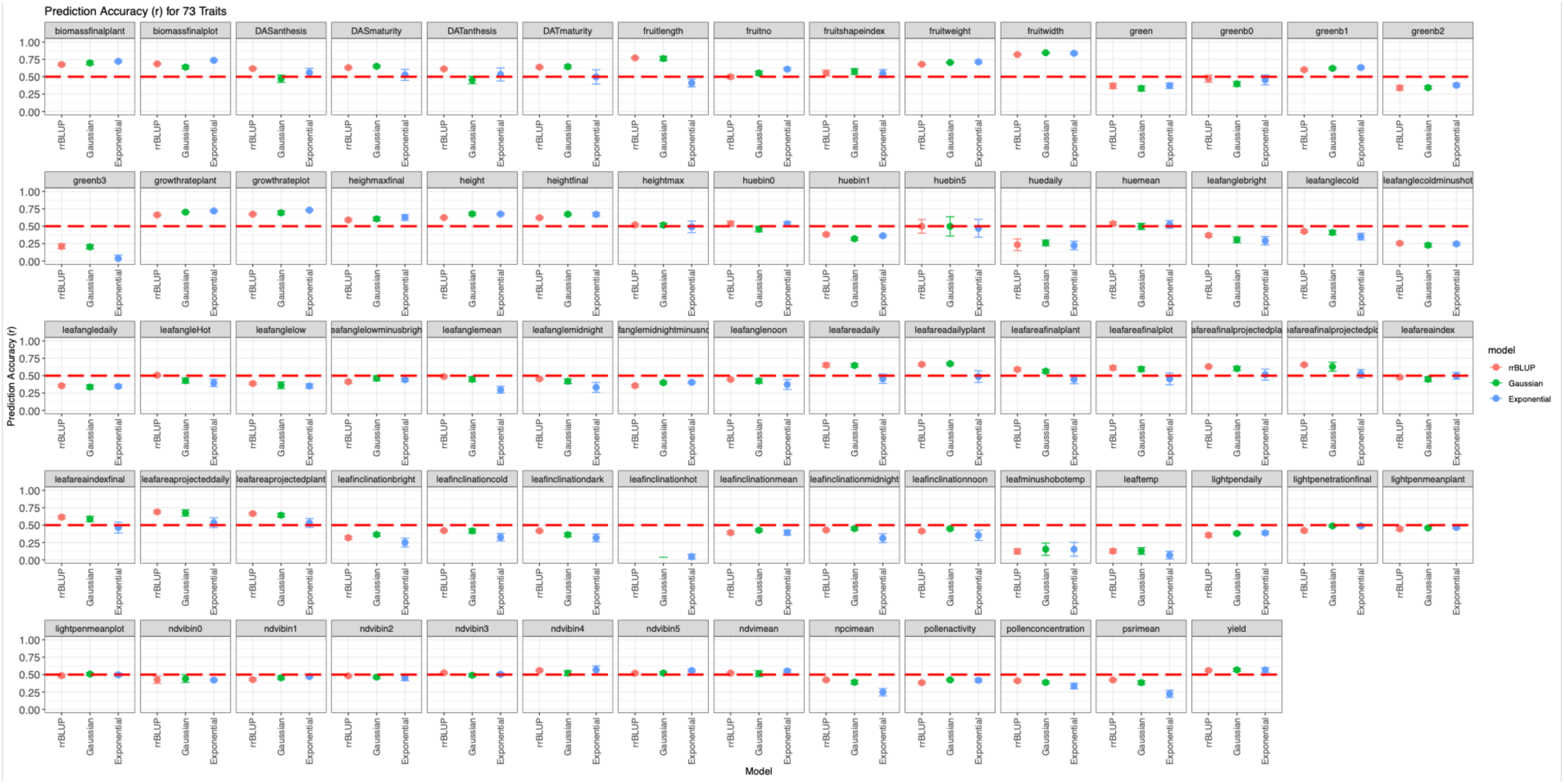
Prediction Accuracies for GS on STI index for 73 agronomic traits for the global *Capsicum* collection. 20 traits were above 0.5 prediction accuracy across all 3 models tested.

## Tables

**Table S1.** 73 phenotypic traits collected at the World Vegetable Centre in Taiwan.

**Table S2.** List of accessions for the core *Capsicum* collection (n = 423) and associated metadata, including principal component (PC) scores for PC1-PC4.

**Table S3.** 23 quality traits from McLeod et al. 2023.

**Table S4.** List of accessions for the global *Capsicum* collection (n = 10,038) and associated metadata. *differs from 10,250 as “CTRL” lines are present in vcf.

**Table S5.** Significant SNPs identified by genome-wide association analysis of stress tolerance index (STI) values across 73 agronomic traits in the *Capsicum annuum* core collection. SNPs passing a Benjamini-Hochberg false discovery rate threshold of FDR < 0.05 in two or more GWAS models (BLINK, FarmCPU, MLM, MLMM) are reported.

**Table S6.** Multi-trait SNPs significant across three or more traits in GWAS of STI values, irrespective of model support.

**Table S7.** High-confidence multi-trait SNPs significant across three or more traits and three or more GWAS models (n = 46), identified from GWAS of STI values in the *Capsicum annuum* core collection.

**Table S8.** PC-adjusted percent variance explained (PVE) for each high-confidence SNP × trait combination.

**Table S9.** GEBVs for the control timepoint in the **core** *Capsicum* collection (n = 423, using 340,734 SNPs and 73 phenotypic traits).

**Table S10.** GEBVs for the heat stress-1 timepoint in the **core** *Capsicum* collection (n = 423, 340,734 SNPs and 73 phenotypic traits).

**Table S11.** GEBVs for the heat stress-2 timepoint in the **core** *Capsicum* collection (n = 423, 340,734 SNPs and 73 phenotypic traits).

**Table S12.** Heat responsive lines (n = 28).

**Table S13.** Control responsive lines (n = 57).

**Table S14.** GEBVs for the control timepoint in the **globa**l *Capsicum* collection (n = 10,250, 23,462 SNPs and 73 phenotypic traits).

**Table S15.** GEBVs for the heat stress-1 timepoint in the **global** *Capsicum* collection (n = 10,250, 23,462 SNPs and 73 phenotypic traits).

**Table S16.** GEBVs for the heat stress-2 timepoint in the **global** *Capsicum* collection (n = 10,250, 23,462 SNPs and 73 phenotypic traits).

**Table S17.** GEBVs for the stress tolerance index (STI) for the **core** *Capsicum* collection (n = 423, 340,734 SNPs and 73 phenotypic traits).

**Table S18.** GEBVs for the stress tolerance index (STI) for the **global** *Capsicum* collection (n = 10,026, 23,462 SNPs and 73 phenotypic traits).

**Table S19.** Genomic selection for 23 quality and traditional traits for the core *Capsicum* collection.

**Table S20.** Genomic selection for 23 quality and traditional traits for the global *Capsicum* collection.

## References

Ambali, M., Zohoungbogbo, H., Ayenan, M., Eybishitz, A., Barchenger, D., & Schreinemachers, P. 2025. Market segmentation for peppers and tomatoes in Africa. Market Intelligence Brief Series 23, Montpellier: CGIAR. https://cgspace.cgiar.org/items/2221b026-cef7-4673-81bb-2fc6f0b53e10

Bates, D., Mächler, M., Bolker, B., & Walker, S. (2015). Fitting Linear Mixed-Effects Models Using lme4. Journal of Statistical Software, 67(1). 10.18637/jss.v067.i01

Borovsky, Y.; Paran, I. Characterization of Fs10.1, a Major QTL Controlling Fruit Elongation in *Capsicum*. Theor. Appl. Genet. 2011, 123, 657–665.

Brown, A. H. D. (1989). Core collections: a practical approach to genetic resources management. Genome, 31(2), 818–824.

Byrne, P. F., Volk, G. M., Gardner, C., Gore, M. A., Simon, P. W., & Smith, S. (2018). Sustaining the future of plant breeding: The critical role of the USDA-ARS National Plant Germplasm System. Crop Science, 58(2), 451–468.

Chaim, A.B.; Borovsky, Y.; Rao, G.U.; Tanyolac, B.; Paran, I. *Fs3.1:* A Major Fruit Shape QTL Conserved in *Capsicum*. Genome 2003, 46, 1–9.

Challinor, A. J., Watson, J., Lobell, D. B., Howden, S. M., Smith, D. R., & Chhetri, N. (2014). A meta-analysis of crop yield under climate change and adaptation. Nature Climate Change, 4(4), 287–291. 10.1038/nclimate2153

Cheng, J.; Qin, C.; Tang, X.; Zhou, H.; Hu, Y.; Zhao, Z.; Cui, J.; Li, B.; Wu, Z.; Yu, J.;, et al. Development of a SNP Array and Its Application to Genetic Mapping and Diversity Assessment in Pepper (Capsicum spp.). Sci. Rep. 2016, 6, 33293.

Cooper, M., Voss-Fels, K. P., Messina, C. D., Tang, T., & Hammer, G. L. (2021). Tackling G × E × M interactions to close on-farm yield-gaps: Creating novel pathways for crop improvement by predicting contributions of genetics and management to crop productivity. Theoretical and Applied Genetics, 134(6), 1625–1644. 10.1007/s00122-021-03812-3.

Covarrubias-Pazaran, G. (2016). Genome-Assisted Prediction of Quantitative Traits Using the R Package sommer. PLOS ONE, 11(6), e0156744. 10.1371/journal.pone.0156744

Crossa, J., Perez, P., Hickey, J., Burgueno, J., Ornella, L., Cerón-Rojas, J., et al. (2014). Genomic prediction in CIMMYT maize and wheat breeding programs. Heredity, 112(1), 48–60.

Danecek, P., Auton, A., Abecasis, G., Albers, C. A., Banks, E., DePristo, M. A., Handsaker, R. E., Lunter, G., Marth, G. T., Sherry, S. T., McVean, G., Durbin, R., & Group, 1000 Genomes Project Analysis. (2011). The variant call format and VCFtools. Bioinformatics, 27(15), 2156–2158. 10.1093/bioinformatics/btr330

Danecek, P., Bonfield, J. K., Liddle, J., Marshall, J., Ohan, V., Pollard, M. O., Whitwham, A., Keane, T., McCarthy, S. A., Davies, R. M., & Li, H. (2021). Twelve years of SAMtools and BCFtools. GigaScience, 10(2). 10.1093/gigascience/giab008

Dempewolf, H., Baute, G., Anderson, J., Kilian, B., Smith, C., & Guarino, L. (2017). Past and future use of wild relatives in crop breeding. Crop science, 57(3), 1070–1082.

DePristo, M. A., Banks, E., Poplin, R., Garimella, K. V, Maguire, J. R., Hartl, C., Philippakis, A. A., del Angel, G., Rivas, M. A., Hanna, M., McKenna, A., Fennell, T. J., Kernytsky, A. M., Sivachenko, A. Y., Cibulskis, K., Gabriel, S. B., Altshuler, D., & Daly, M. J. (2011). A framework for variation discovery and genotyping using next-generation DNA sequencing data. Nature Genetics, 43(5), 491–498. 10.1038/ng.806

Desta, Z. A., & Ortiz, R. (2014). Genomic selection: genome-wide prediction in plant improvement. Trends in plant science, 19(9), 592–601.

DeWitt, D., & Bosland, P. W. (1993). The pepper garden. Berkeley, CA: Ten Speed Press.

Escamilla, D. M., Li, D., Negus, K. L., Kappelmann, K. L., Kusmec, A., Vanous, A. E., et al. (2025). Genomic selection: Essence, applications, and prospects. The Plant Genome, 18(2), e70053.

Fernandez, G.C.J. (1992) Effective Selection Criteria for Assessing Stress Tolerance. In: Kuo, C.G., Ed., Proceedings of the International Symposium on Adaptation of Vegetables and Other Food Crops in Temperature and Water Stress, AVRDC Publication, Tainan, 257–270.

Finlay, K., & Wilkinson, G. (1963). The analysis of adaptation in a plant-breeding programme. Australian Journal of Agricultural Research, 14(6), 742–754. 10.1071/AR9630742

Guzman, I., Bosland, P. W., & O’Connell, M. A. (2011). Heat, color, and flavor compounds in Capsicum fruit. In D. R. Gang (Ed.), The biological activity of phytochemicals (pp. 109–126). New York, NY: Springer.

Hajjar, R., & Hodgkin, T. (2007). The use of wild relatives in crop improvement: a survey of developments over the last 20 years. Euphytica, 156, 1–13.

Heslot, N., Yang, H. P., Sorrells, M. E., & Jannink, J. L. (2012). Genomic selection in plant breeding: a comparison of models. Crop science, 52(1), 146–160.

Kantar, M. B., Anderson, J. E., Lucht, S. A., Mercer, K., Bernau, V., Case, K. A., et al. (2016). Vitamin variation in Capsicum spp. provides opportunities to improve nutritional value of human diets. PLoS ONE, 11, e0161464. 10.1371/journal.pone.0161464

Khaitov, B., Umurzokov, M., Cho, K.-M., Lee, Y.-J., Park, K. W., & Sung, J. (2019). Importance and production of chilli pepper; heat tolerance and efficient nutrient use under climate change conditions. Korean Journal of Agricultural Science, 46(4), 769–779. 10.7744/kjoas.20190059

Knaus, B. J., & Grünwald, N. J. (2017). vcfr: A package to manipulate and visualize variant call format data in R. Molecular Ecology Resources, 17(1), 44–53. 10.1111/1755-0998.12549

Langridge, P., & Fleury, D. (2011). Making the most of ‘omics’ for crop breeding. Trends in Biotechnology, 29(1), 33–40. 10.1016/j.tibtech.2010.09.006

Lee, S. G., Kim, S. K., Lee, H. J., Lee, H. S., & Lee, J. H. (2018). Impact of moderate and extreme climate change scenarios on growth, morphological features, photosynthesis, and fruit production of hot pepper. Ecology and evolution, 8(1), 197–206.

Lenth R, Piaskowski J (2025). emmeans: Estimated Marginal Means, aka Least-Squares Means. R package version 2.0.1, https://rvlenth.github.io/emmeans/.

Levins, R. (1967). THEORY OF FITNESS IN A HETEROGENEOUS ENVIRONMENT. VI. THE ADAPTIVE SIGNIFICANCE OF MUTATION. Genetics, 56(1), 163–178. 10.1093/genetics/56.1.163

Li, H., Rasheed, A., Hickey, L. T., & He, Z. (2018). Fast-forwarding genetic gain. Trends in plant science, 23(3), 184–186.

Liu, F., Zhao, J., Sun, H., Xiong, C., Sun, X., Wang, X., Wang, Z., Jarret, R., Wang, J., Tang, B., Xu, H., Hu, B., Suo, H., Yang, B., Ou, L., Li, X., Zhou, S., Yang, S., Liu, Z., … Zou, X. (2023). Genomes of cultivated and wild Capsicum species provide insights into pepper domestication and population differentiation. Nature Communications, 14(1), 5487. 10.1038/s41467-023-41251-4

Marklein, A., Elias, E., Nico, P., & Steenwerth, K. (2020). Projected temperature increases may require shifts in the growing season of cool-season crops and the growing locations of warm-season crops. Science of The Total Environment, 746, 140918.

Martínez-Ainsworth, N. E., Scheppler, H., Moreno-Letelier, A., Bernau, V., Kantar, M. B., Mercer, K. L., & Jardón-Barbolla, L. (2023). Fluctuation of ecological niches and geographic range shifts along chile pepper’s domestication gradient. Ecology and Evolution, 13(11), e10731.

McCouch, S., Baute, G. J., Bradeen, J., Bramel, P., Bretting, P. K., Buckler, E., Burke, J. M., Charest, D., Cloutier, S., Cole, G., Dempewolf, H., Dingkuhn, M., Feuillet, C., Gepts, P., Grattapaglia, D., Guarino, L., Jackson, S., Knapp, S., Langridge, P., … Zamir, D. (2013). Feeding the future. Nature, 499(7456), 23–24. 10.1038/499023a

McLeod, L., Barchi, L., Tumino, G., Tripodi, P., Salinier, J., Gros, C., et al. (2023). Multi-environment association study highlights candidate genes for robust agronomic quantitative trait loci in a novel worldwide Capsicum core collection. The Plant Journal, 116(5), 1508–1528.

Moscone, E.A.; Scaldaferro, M.A.; Grabiele, M.; Cecchini, N.M.; García, Y.S.; Jarret, R.; Daviña, J.R.; Ducasse, D.A.; Barboza, G.E.; Ehrendorfer, F. The Evolution of Chili Peppers (Capsicum-Solanaceae): A Cytogenetic Perspective. Acta Hortic. 2007, 745, 137–170.

Olatunji, T. L., & Afolayan, A. J. (2018). The suitability of chili pepper ( *Capsicum annuum* L.) for alleviating human micronutrient dietary deficiencies: A review. Food Science & Nutrition, 6(8), 2239–2251. 10.1002/fsn3.790

Ortiz-Bobea A, Ault TR, Carrillo CM, Chambers RG, Lobell DB. 2021. Anthropogenic climate change has slowed global agricultural productivity growth. Nat Clim Chang. 11(4):306–312. 10.1038/s41558-021-01000-1.

Purcell, S., Neale, B., Todd-Brown, K., Thomas, L., Ferreira, M. A. R., Bender, D., Maller, J., Sklar, P., de Bakker, P. I. W., Daly, M. J., & Sham, P. C. (2007). PLINK: A Tool Set for Whole-Genome Association and Population-Based Linkage Analyses. The American Journal of Human Genetics, 81(3), 559–575. 10.1086/519795

Paran, I.; van der Knaap, E. Genetic and Molecular Regulation of Fruit and Plant Domestication Traits in Tomato and Pepper. J. Exp. Bot. 2007, 58, 3841–3852.

Ren, R., Zhou, X., & Feng, J. (2024). Wheat disease resistance: diagnosis, germplasm mining, and molecular breeding. Frontiers in Plant Science, 15, 1500414.

Ro, N., Oh, H., Ko, H. C., Yi, J., Na, Y. W., & Haile, M. (2024). Exploring Genomic Regions Associated with Fruit Traits in Pepper: Insights from Multiple GWAS Models. International Journal of Molecular Sciences, 25(21), 11836.

Sadrarhami, A., Khoshgoftarmanesh, A. H., & Sharifi, H. R. (2010). Using stress tolerance indicator (STI) to select high grain yield iron-deficiency tolerant wheat genotypes in calcareous soils. Field Crops Research, 119(1), 12–19.

Sanchez, D., Sadoun, S. B., Mary-Huard, T., Allier, A., Moreau, L., & Charcosset, A. (2023). Improving the use of plant genetic resources to sustain breeding programs’ efficiency. Proceedings of the National Academy of Sciences, 120(14), e2205780119.

Schrauf, M. F., de Los Campos, G., & Munilla, S. (2021). Comparing genomic prediction models by means of cross validation. Frontiers in Plant Science, 12, 734512.

Sunitha, N. C., Prathibha, M. D., Thribhuvan, R., Lokeshkumar, B. M., Basavaraj, P. S., Lohithaswa, H. C., & Anilkumar, C. J. G. R. (2024). Focused identification of germplasm strategy (FIGS): a strategic approach for trait-enhanced pre-breeding. Genetic Resources and Crop Evolution, 71(1), 1–16.

Tigchelaar, M., Battisti, D. S., Naylor, R. L., & Ray, D. K. (2018). Future warming increases probability of globally synchronized maize production shocks. Proceedings of the National Academy of Sciences, 115(26), 6644–6649. 10.1073/pnas.1718031115

Tigchelaar, M., Selig, E. R., Sarhadi, A., Bruce, J., Allison, E. H., Battista, W., et al. (2024). Nutrition-sensitive climate risk across food production systems. Environmental Research Letters, 20(1), 014046.

Wall, M. M., & Bosland, P. W. (1998). Analytical methods for color and pungency of chiles (Capsicums). In D. Wetzel, & G. Charalambous (Eds.), Instrumental methods in food and beverage analysis (pp. 347–373). Amsterdam, The Netherlands: Elsevier B.V.

Wambugu, P. W., Ndjiondjop, M. N., & Henry, R. J. (2018). Role of genomics in promoting the utilization of plant genetic resources in genebanks. Briefings in functional genomics, 17(3), 198–206.

Wang, J., & Zhang, Z. (2021). GAPIT version 3: boosting power and accuracy for genomic association and prediction. Genomics, proteomics & bioinformatics, 19(4), 629–640.

Yang, J., Sears, R. G., Gill, B. S., & Paulsen, G. M. (2002). Quantitative and molecular characterization of heat tolerance in hexaploid wheat. Euphytica, 126(2), 275–282.

Yu, X., Li, X., Guo, T., Zhu, C., Wu, Y., Mitchell, S. E., et al. (2016). Genomic prediction contributing to a promising global strategy to turbocharge gene banks. Nature plants, 2(10), 1–7.

Yu, X., Leiboff, S., Li, X., Guo, T., Ronning, N., Zhang, X., et al. (2020). Genomic prediction of maize microphenotypes provides insights for optimizing selection and mining diversity. Plant Biotechnology Journal, 18(12), 2456–2465.

Zhang, H., Zhu, J., Gong, Z., & Zhu, J. K. (2022). Abiotic stress responses in plants. Nature Reviews Genetics, 23(2), 104–119.

